# Periplasmic Expression of SpyTagged Antibody Fragments Enables Rapid Modular Antibody Assembly

**DOI:** 10.1101/2020.09.18.302950

**Authors:** Christian Hentrich, Sarah-Jane Kellmann, Mateusz Putyrski, Manuel Cavada, Hanh Hanuschka, Achim Knappik, Francisco Ylera

**Affiliations:** Bio-Rad AbD Serotec GmbH, Zeppelinstraße 4, 82178 Puchheim, Germany

## Abstract

Antibodies are essential tools in research and diagnostics. While antibody fragments can be rapidly produced in *Escherichia coli*, full-length antibodies with an Fc region or antibodies modified with probes are time and labor intensive in production.

SpyTag/SpyCatcher protein ligation technology could covalently attach such functionalities to antibody fragments equipped with a SpyTag. However, we found that the necessarily periplasmic expression of such antibody fragments in *E. coli* led to rapid cleavage of the SpyTag by proteases.

Here we show how this cleavage can be prevented, making the SpyTag technology accessible for *E. coli* produced antibodies. We demonstrate a modular toolbox for rapid creation of synthetic IgGs, oligomerized antibodies, and antibodies with different tags or enzymatic functionalities and measure their performance in a variety of immunoassays. Furthermore, we demonstrate surface immobilization, high-throughput screening of antibody libraries, and rapid prototyping of antibodies based on modular antibody assembly.

## Introduction

Antibodies are valuable tools for research and development in the life sciences and for medical diagnostics^1, 2^. They are used in many different applications and, with the rise of antibody engineering in the last decades, many recombinant antibody formats like Fab fragments (fragment antigen-binding, Fabs), single-chain variable fragments (scFv), diabodies, or nanobodies became available^3^. As each experiment utilizing antibodies has different requirements, it is beneficial to use an antibody format optimized for the application. For example, for the measurement of antibody affinities, monovalent antibody formats such as Fabs are advantageous by avoiding confounding avidity effects. On the other hand, avidity is highly desirable in many assays such as immunoblotting or enzyme-linked immunosorbent assay (ELISA) to increase the overall sensitivity and therefore full-length immunoglobulins or dimeric Fabs (e.g. F(ab’)_2_) are the preferred antibody format. Similar optimizations can be made by changing the Fc part dependent on the secondary anti-Fc reagents to be used, or by adding dyes or labels to the antibody prior to performing the assay.

Traditionally, full immunoglobulin G (IgG) antibodies are most commonly used for assays because IgG is the main isotype generated by the host’s immune system and the default product of monoclonal antibody production in hybridoma cell lines^4^. The Fc part of IgGs is often solely required for detection with a secondary reagent and is not a core component of the assay. Therefore, in many assays, IgGs can be replaced with bacterially expressed antibody fragments such as Fabs or scFvs, which can be produced much more economically. Functional antibody fragment expression in *E. coli* is typically achieved by using the bacterial transport apparatus to direct the nascent antibody chains into the oxidizing environment of the periplasm, where the disulfide bonds necessary for antibody folding can form^5, 6^.

In an assay, detection of the primary antibody is most often performed with polyclonal secondary antibodies raised in another species and labeled with a suitable probe or an enzyme. This avoids the laborious and expensive labeling of each primary antibody. However, use of secondary antibodies might introduce unspecific binding which needs to be controlled and, furthermore, extends the duration of the assay by additional incubation and washing steps^7^. That is why, especially for routine assays, labeled primary antibodies are preferred. They are also useful for multiplex assays, especially in flow cytometry, with each primary antibody directly labeled with a different fluorescent dye. It stands to reason that directly labeled antibodies would be the preferred format in many assays if they were easier to generate or were commercially available.

Labeling of antibodies with enzymes or probes is most commonly carried out using amine-reactive functional groups such N-hydroxysuccinimide (NHS), or Traut’s reagent in combination with maleimide, which are not site-specific^8^. This results in high batch-to-batch variability and the risk of modifying the antigen binding site and thus impairing the function of the antibody. To circumvent this, methods have been developed to target specific chemical groups on the Fc chain such as the sugar moieties or the Fc disulfide bonds^8, 9^. Common to most chemical labeling strategies is that they often require high protein concentrations, long incubation times, or additional purification steps.

Proteins can be site-specifically labeled through protein ligation, in which a covalent bond is formed between two polypeptide chains either autocatalytically or through an external enzyme. Several methods have been developed in the past decades, such as sortase-mediated ligation or intein-based approaches^10^. A recent advance was the development of split proteins from proteins that naturally contain an isopeptide-bond between amino acid side chains. When mixed together, the split protein segments re-associate and autocatalytically form the isopeptide bond contained in the wildtype protein, thus covalently crosslinking two independent polypeptides. The most prominent of these technologies is the SpyTag/SpyCatcher system which was developed through protein engineering from the fibronectin binding protein FbaB from *Streptococcus pyogenes*. In this system, SpyTag, a small peptide of 13 amino acids, reacts spontaneously with SpyCatcher, a small protein of 12.3 kDa, to form an isopeptide bond^11^ (Figure 1A). The reaction is fast (within minutes to an hour), specific, and several optimized versions of the reaction partners have been developed, termed SpyTag2/SpyCatcher2^12^ and SpyTag3/SpyCatcher3^13^. For clarity, we will be using the terms SpyTag1/SpyCatcher1 when referring specifically to the original SpyTag/SpyCatcher protein sequences and will use the unnumbered names as generic terms. The versatility of this system was shown in many publications in a broad range of different applications and has been the subject of several review articles^14, 15^.

**Figure 1:**
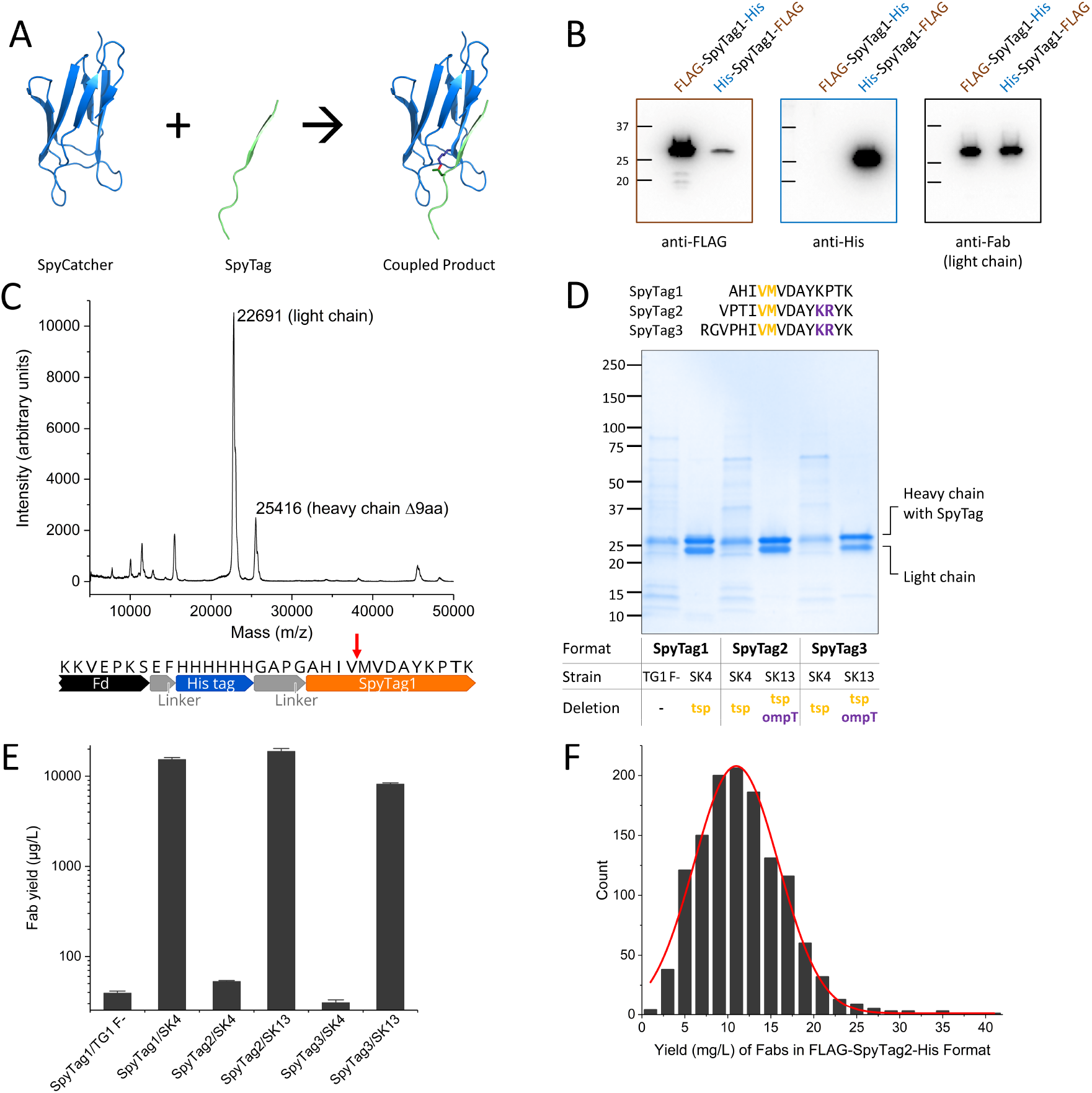
Expression of SpyTagged Fabs in the Periplasm of *E. coli*. **A**: SpyCatcher (blue) reacts with SpyTag (green) to form a covalent isopeptide bond (red). Images generated from PDB Structure 4MLI^50^. **B**: Immunoblots of crude bacterial lysates from overnight expression cultures of a human Fab in FLAG-Spy-His and His-Spy-FLAG format. Blots were probed with HRP-labeled anti-FLAG (left), anti-His (middle), or anti-Fab (right). Molecular weights are indicated in kDa. **C**: Linear mode MALDI Mass spectrum (top) of a Ni-NTA affinity-purified Fab in His-SpyTag1 format (bottom). Expected protonated full-length masses: 22,691 Da (light chain), 26,437 Da (heavy chain). Expected protonated mass for the heavy chain with a 9 amino acid C-terminal truncation: 25,403 Da. The putative cleavage site is indicated by a red arrow. **D**: Top: Amino acid sequences of SpyTag1/2/3 with the amino acids surrounding the putative protease cleavage sites highlighted in color. Bottom: SDS-PAGE of Fabs equipped with different SpyTag formats and purified from different bacterial strains. 2 µg of total protein as determined by A280 measurement were loaded in each lane. Molecular weights are indicated in kDa. **E**: Concentration of SpyTagged Fabs after purification from different bacterial strains as determined by titration ELISA (capture with anti-Fab, detection with anti-His-HRP). **F**: Histogram of the yield per liter of *E. coli* culture from 1,281 purifications of Fabs in FLAG-SpyTag2-His format in the SK13 double knockout strain.

We set out to develop a generic way to completely bypass laborious cloning steps for the generation of different antibody formats, taking advantage of the SpyTag technology to assemble recombinant antibodies in various forms and with a number of labels by pre-fabricated protein modules. We envisioned bacterially produced Fabs equipped with a SpyTag that could be coupled to a diverse array of independently produced SpyCatcher variants, in different oligomeric states, and equipped with different functionalities.

Proof of concept for such an approach was demonstrated for one antibody by Geyer and colleagues^16^, and two publications^17, 18^ describe periplasmic bacterial expression of nanobodies with a SpyTag. However, we observed that the vast majority of SpyTagged antibodies cannot be produced in functional form in the periplasm due to a pronounced instability of the SpyTag in the periplasm of *E. coli*.

In this study, we identified the proteases responsible for SpyTag cleavage, and we present strategies to address this instability and to enable robust access to periplasmically expressed SpyTag fusion proteins. We then present a toolbox of SpyCatcher-based adapter proteins which can be coupled quantitatively to the antibodies within one hour and we demonstrate the performance of these coupled products in a range of typical antibody assays. Finally, we show applications in which the SpyTag/SpyCatcher system not only enables improvements in convenience or assay sensitivity, but also allows the development of novel screening concepts for *in vitro* selections and improved robustness in assay design.

## Results

### Degradation of the SpyTag in the Periplasm

Human Fab antibody fragments are routinely expressed in the periplasm of *E. coli*, in which oxidizing conditions allow the formation of disulfide bonds needed for stable folding^5^. In order to allow site-specific labeling of Fabs via the SpyTag technology, we constructed plasmids for periplasmic expression of human Fabs where the heavy chain was fused to a C-terminal SpyTag1 followed by a hexahistidine tag (His-tag). To our surprise, we were unable to purify antibodies with an intact SpyTag1 from *E. coli* in this manner. We hypothesized that the SpyTag1 might be cleaved in the bacterial periplasm, thereby removing the C-terminal His-tag required for Fab purification. To test this, we constructed Fabs in which the heavy chain was C-terminally tagged with a SpyTag1 encompassed between a FLAG-tag and a His-tag in both orientations (i.e. Fab-FLAG-Spy-His and Fab-His-Spy-FLAG). We performed immunoblots against the FLAG-tag and His-tag in crude bacterial lysates after overnight expression of the aforementioned constructs (Figure 1B). In the Fab-FLAG-SpyTag1-His configuration, we could detect a strong band corresponding to the size of the heavy chain in the anti-FLAG blot, but not in the anti-His blot. For the inverse tag order (Fab-His-SpyTag1-FLAG), we obtained inverse immunoblotting results (Figure 1B). This observation suggested that the SpyTag was indeed cleaved during overnight expression without affecting expression of the Fab part of the construct. To characterize this further, we constructed a Fab with the heavy chain fused to a C-terminal His-tag followed by a SpyTag1 and expressed it overnight in *E. coli*. We purified the protein via Ni-NTA affinity chromatography and examined the resulting protein via linear mode matrix-assisted laser desorption/ionization time-of-flight (MALDI-TOF) mass spectrometry (Figure 1C). The expected protonated mass of the light chain was detected precisely, however there was no peak for the full-length heavy chain mass. Instead, we found a peak corresponding to a heavy chain truncated by 9 amino acids, consistent with a cleavage of the SpyTag1 after the first valine residue. Further support for a truncation at the C-terminus was provided by peptide mass fingerprint MALDI TOF mass spectrometry, in which the very N-terminal peptide could be detected, indicating that cleavage occurred at the C-terminus instead (Supplementary figure S1). We performed equivalent tests with a SpyTagged maltose binding protein (MBP) expressed in the periplasm, and again observed cleavage of the SpyTag1 (Supplementary figure S2), demonstrating the SpyTag1 instability is not limited to antibody fragments. The hypothesis that the cleavage occurred in the periplasm was supported by the fact that full-length SpyTagged MBP could be expressed in bacterial cytoplasm without any difficulty.

### Enabling Periplasmic Expression of SpyTagged Fab Fragments

As these results hinted at proteolytic cleavage, we focused our attention on proteases active in the bacterial periplasm, especially those that were described to cleave after a valine at the C-terminus of a protein. Tail-specific protease (Tsp, also termed Prc) has been described to cleave after nonpolar residues at the C-terminus of proteins^19^ and to reside on the periplasmic side of the bacterial cell membrane^20^.

To test the influence of Tsp on SpyTag1 expression, we constructed a tsp knock out variant of our *E. coli* expression strain TG1 F-using the method of Datsenko and Wanner^21^ (Supplementary figure S3). The TG1 F-Δtsp knockout cells, termed SK4, did not exhibit any obvious growth phenotype. Strikingly, when we repeated the expression of our SpyTagged Fab constructs in SK4 cells, the SpyTag1 remained intact (Figure 1D, lane2) and Fabs could be purified with good yields (Figure 1E).

When we attempted to express SpyTag2^12^, the first improved version of SpyTag1, in SK4 cells, we again could not purify full-length Fab-FLAG-SpyTag2-His (Figure 1E). We repeated the mass spectrometry experiment with a Fab heavy chain with a C-terminal His-tag followed by a SpyTag2, expressed in SK4 cells, and detected a potential proteolytic cleavage site within the last four amino acids of SpyTag2 (Supplementary figure S4). Outer membrane protein T (OmpT) is a protease active in the periplasm with a recognition motif centered around two consecutive positively charged amino acids^22^. Such a motif is present at the C-terminus of SpyTag2 (…Y**KR**YK), but not SpyTag1 (…YKPTK). We therefore proceeded to generate the double knockout *E. coli* strain TG1 F-Δtsp ΔompT, termed SK13 (Supplementary figure S3). This double knockout strain exhibited mild growth defects when cultured at 37°C in rich medium, but this did not affect its usability as an expression strain at lower growth temperatures. Importantly, the SK13 strain did indeed enable expression of Fabs with a full-length SpyTag2 (Figure 1D) with good yields (Figure 1E). Fabs fused to SpyTag3, which shares the putative ompT cleavage site with SpyTag2, could also be successfully produced in SK13 double knockout cells, but not in SK4 single knockout (Figure 1D, E).

Before the knockout strains became available, we developed an alternative method for periplasmic SpyTag expression. We reasoned that it might be also possible to prevent proteolytic cleavage by sterically blocking Tsp protease from accessing the SpyTag substrate. Indeed, moving the SpyTag1 close to the C-terminus of the Fab heavy chain by deleting the linker sequence protected SpyTag1 from proteolysis by Tsp protease while retaining functionality for coupling with SpyCatcher (Supplementary figure S5A, B). Proximity to a folded domain impaired reaction speed of coupling, and the best optimal combination of reactivity and proteolytic stability was found to be a linker sequence of 4 hinge residues (Supplementary figure S5C, D). This strategy however did not protect against cleavage of the SpyTag2 by ompT, since this protease cleaves at the end of the tag and therefore protease access cannot be hindered by proximity to a folded domain.

In summary, we have shown that all variants of the SpyTag are cleaved by periplasmic proteases and developed strategies to mitigate this problem. Interestingly, Alam *et al*. have reported successful Fab-SpyTag1 expression in the periplasm for one antibody^16^ without using these approaches. We reproduced the precise antibody format reported in this publication with 12 different antibodies and measured abundance of full-length Fab antibody when expressed in TG1 F- or BL21 *E. coli* strains (the latter was used in Alam et al.), relative to abundance when expressed in the SK4 strain (Supplementary figure S6). The amount of full-length Fab from SK4 cells was more than 10-fold higher in all cases.

For further experiments, we chose the Fab-FLAG-SpyTag2-His format (short: F-Spy2-H) as it showed clearly improved coupling speed over SpyTag1. We expressed over 1,000 different human Fab antibodies in this format, with a median yield of 11 mg per liter of *E. coli* culture (Figure 1F), very similar to the yield reported for HuCAL Fab antibodies without SpyTag^23^.

### A Modular Antibody Assembly Toolbox

Next, we set out to generate a diverse portfolio of SpyCatcher molecules. In SpyCatcher2 we noticed frequent deamidation at asparagine 105, and mutated it to aspartate, the same amino acid in place in SpyCatcher3 (Supplementary figure S7). We generated SpyCatcher2, BiCatcher2 (a homodimer of SpyCatcher2 bridged with a flexible linker), FcCatcher3 (SpyCatcher3 fused to the Fc region of various immunoglobulin isotypes from different species), and MultiCatchers (higher-order SpyCatcher2 oligomers fused with flexible linkers). All SpyCatcher2 variants carried the N105D mutation except for the MultiCatchers. Furthermore, we produced variants containing unpaired cysteines for site-specific labeling with maleimide chemistry, allowing us to produce SpyCatchers labeled with horseradish peroxidase (HRP) or phycoerythrin (PE) (Figure 2A). Similar constructs based on SpyCatcher1 have been described in the literature^16, 24, 25^.

**Figure 2:**
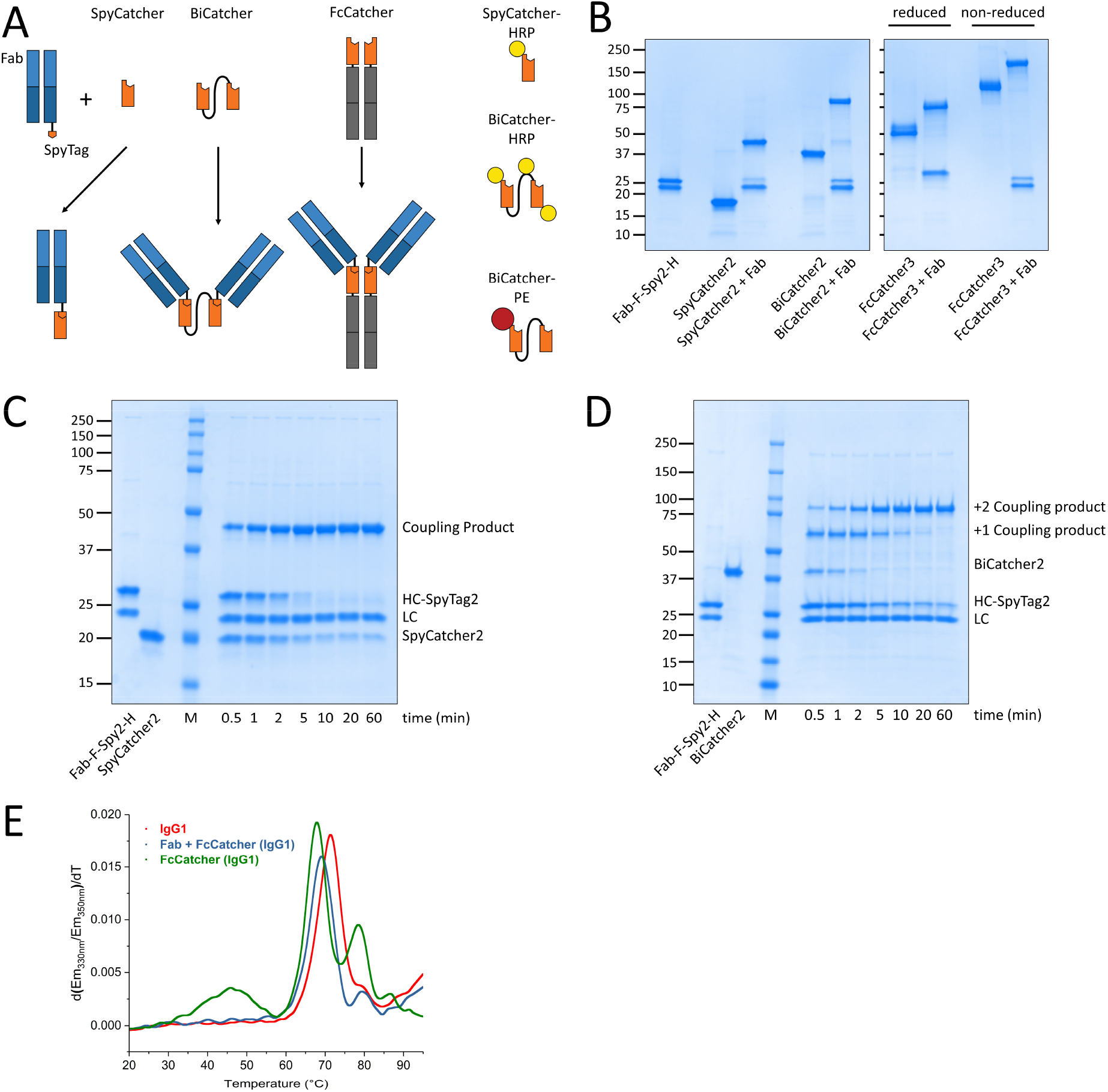
SpyCatcher-based Modular Building Blocks. **A**: Scheme of modular antibody construction using SpyTagged Fabs and SpyCatcher derived building blocks. Fabs (blue) with a SpyTag (orange pentagon) can react with SpyCatcher (orange polygon), to form a monomeric functionalized Fab. Alternatively, dimeric forms can be generated by reacting the Fab with BiCatcher, generating a dimeric Fab, or FcCatcher (grey: Fc), generating an IgG-like protein. SpyCatcher and BiCatcher can be site-specifically labeled with functionalities like HRP (yellow) or PE (red). **B**: SDS-PAGE of building blocks uncoupled and coupled to a SpyTagged Fab. Fabs are in 25% molar excess. Molecular weights are indicated in kDa. **C**: SDS-PAGE time course of the coupling reaction of 4 µM Fab-SpyTag2 and 5 µM SpyCatcher2. Molecular weights are indicated in kDa, the molecular weight marker lane is labeled ‘M’. **D**: SDS-PAGE time course of the coupling reaction of 10 µM Fab-SpyTag2 and 4 µM BiCatcher. Molecular weights are indicated in kDa, the molecular weight marker lane is labeled ‘M’ **E**: NanoDSF curves of FcCatcher3 (hIgG1_FcSC3) alone or coupled to a Fab in comparison with the equivalent IgG. Measurements were performed at approximately 0.4 mg/mL total protein concentration.

As expected, the Fab-SpyTag fusion proteins reacted readily with the various SpyCatcher proteins (Figure 2B). Reactions with monomeric SpyCatcher2 were completed after ten minutes (Figure 2C), whereas dimeric constructs need longer reaction times, up to one hour, possibly because of steric hindrance during the second coupling step (Figure 2D). Particularly for these constructs, the improved versions of SpyCatcher are helpful to keep the reaction time short. To further characterize the coupling reaction, we measured melting curves of uncoupled and coupled FcCatchers via nanoscale differential scanning fluorimetry^26^ and compared them to the analogous wildtype full-length immunoglobulins. For uncoupled FcCatcher, a first melting point of 45°C could be detected (Figure 2E), similar to isolated SpyCatcher2 (Supplementary figure S8). FcCatcher coupled to Fab-FLAG-SpyTag2-His showed a first melting point at 68°C, very similar to the wildtype IgG1 (Figure 2E). This stabilization of the SpyCatcher domain by covalently bound SpyTag is also observed with isolated SpyCatcher2 and other FcCatcher isotypes (Supplementary figure S8).

### Assay Performance

Next, we tested how the diverse Fab-SpyTag-SpyCatcher coupled products perform in typical antibody-based assays in comparison to conventional antibodies. In immunoblotting, we could observe that higher avidity through dimeric or higher-order MultiCatchers led to increased sensitivity compared to monomeric Fabs (Figure 3A). We could also observe that BiCatcher labeled with three copies of HRP via engineered cysteines led to higher assay sensitivity than detection with commercially available polyclonal secondary antibodies (Figure 3B). While performing immunoblotting experiments, we noticed that uncoupled SpyCatcher-HRP gave a weak but specific signal with a band of approximately 51 kDa, using lysates of several eukaryotic cell lines. We were not able to identify the protein corresponding to this band, but we established that the appearance of the band could be avoided via full coupling of the SpyCatcher-HRP by adding sufficient Fab-SpyTag during the coupling reaction, or through quenching the reaction by addition of an excess of free SpyTag3 peptide (Supplementary figure S9).

**Figure 3:**
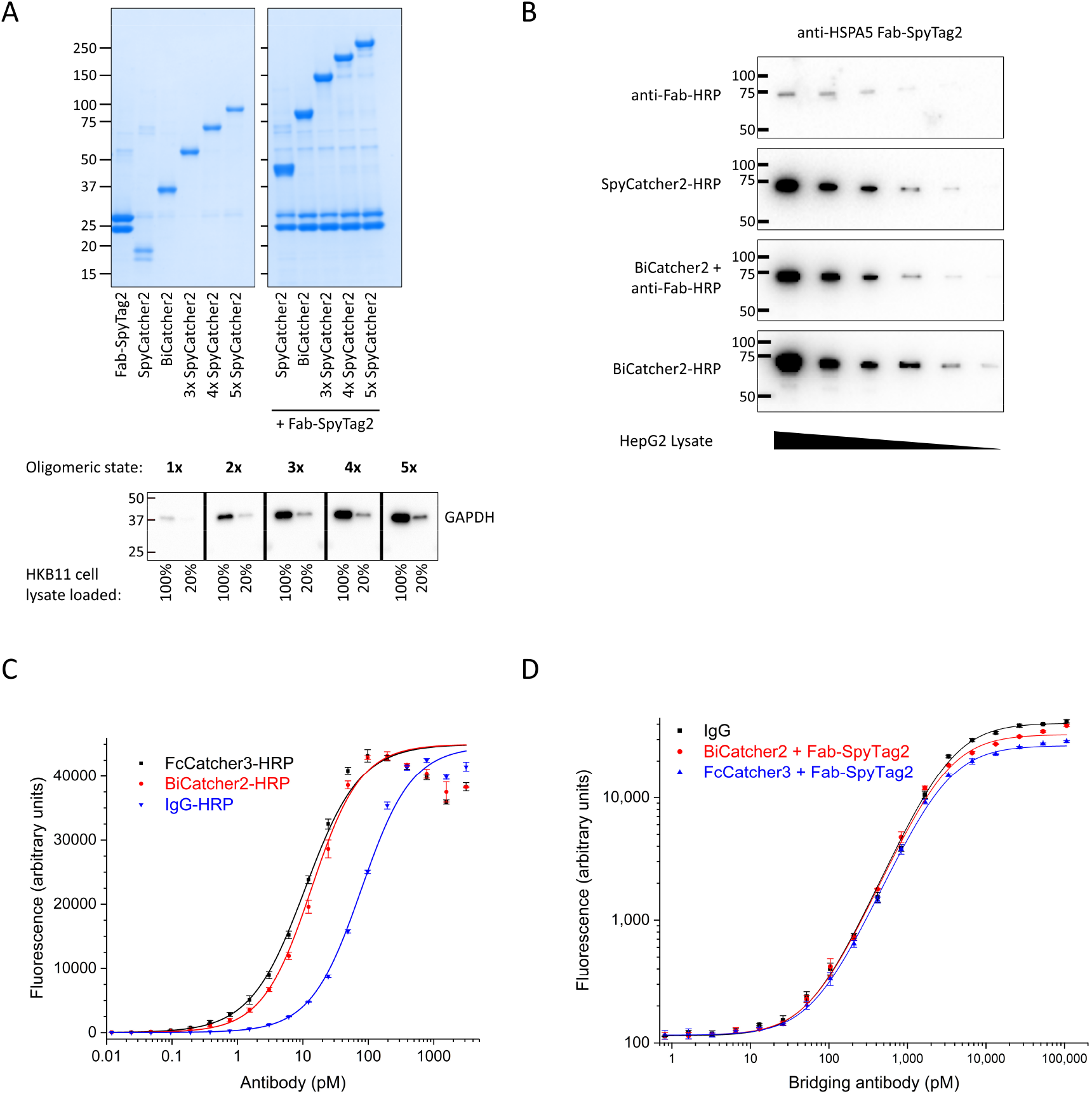
Performance of Assembled Modular Antibodies in Immunoblotting and ELISA. **A**: Top: SDS PAGE of SpyCatcher oligomerized from 1x to 5x, and of coupling reactions with SpyTagged anti-GAPDH Fab (clone AbD36683). Bottom: Immunoblots of HKB11 cell lysates loaded at two dilutions probed with the mono- and oligomeric Fab-SpyCatcher fusion proteins shown on the left. Detection was performed with anti-Fab-HRP. Molecular weights are indicated in kDa. **B**: Immunoblots of HepG2 total cell lysate (Bio-Rad) loaded in a 1:2 dilution series probed with anti-HSPA5 Fab-FLAG-SpyTag2-His antibody (clone AbD23588ad). Detection was performed via anti-Fab-HRP antibody (Bio-Rad) as a secondary reagent, via direct detection with Fab coupled to SpyCatcher2-HRP, with Fab coupled to BiCatcher2 and anti-Fab-HRP antibody as a secondary reagent, or via direct detection with Fab coupled to BiCatcher2-HRP. **C**: Titration ELISA of anti-trastuzumab antibody (clone AbD18018) as HRP-labeled IgG or as Fab-FLAG-SpyTag2-His coupled to BiCatcher2-HRP or FcCatcher3-HRP. Trastuzumab is coated at 1 µg/mL. Fits are logistic and do not fit the hook effect observed with HRP-labeled SpyCatchers. **D**: Bridging ELISA with an anti-ipilimumab antibody (clone AbD34428) as IgG or as Fab-FLAG-SpyTag2-His coupled to BiCatcher2 or FcCatcher3. Ipilimumab is coated at 1 µg/mL, detection is performed with HRP-labeled ipilimumab at 2 µg/mL. Error bars are standard deviations of measurements in triplicates.

Performance of Fab antibodies labeled with SpyCatcher-HRP was then compared with conventionally HRP-labeled IgG in ELISA (Figure 3C). In direct titration ELISA, HRP-labeled BiCatcher2 or FcCatcher3 showed better sensitivity than IgGs labeled via an HRP conjugation kit (LYNX, Bio-Rad). The dimeric nature of BiCatcher and FcCatcher molecules also allows the use of their coupled products in bridging sandwich ELISA assays, in which one Fab domain binds to a capture antibody, and the other is bound by a detection antibody. Here, using an anti-drug antibody assay set-up as an example, the conjugation products perform similarly to IgGs (Figure 3D), with small differences probably due to altered geometry.

It has been shown that surface-coated SpyCatcher1 can be used as a capture reagent for *in vitro* assays^17, 27^ (Figure 4A). We wanted to establish such an oriented immobilization with the faster-reacting SpyTag2 and SpyCatcher3, which should provide an advantage at the low protein concentrations used for immobilization in ELISA assays. Therefore, we coated polystyrene plates (Maxisorp, Thermo Fisher) with SpyCatcher3 and assayed the coupling kinetics of purified SpyTagged Fabs in serial dilutions by ELISA (Figure 4B). As expected, the coupling was concentration and time dependent, with high antibody concentrations (40 µg/mL) allowing for full coupling already after 5 minutes, whereas concentrations below 1 µg/mL could be effectively coupled within 1-3 hours. It has been reported that oriented immobilization of capture antibodies, where antibodies are not directly coated on surface but immobilized by e.g. another antibody, can be beneficial for assay sensitivity, probably by ensuring that a higher fraction of the antibodies is functional^28^. To test if oriented immobilization through SpyCatcher could also increase the amount of active antibody on the surface, we prepared a dilution series of SpyTagged Fabs and either coupled them to immobilized SpyCatcher (coated at varying concentrations) or coated them directly on the surface and assayed the antigen binding capability (Figure 4C). We observed that the overall amount of active antibody (as indicated by the maximal signal reached) on the surface cannot be substantially increased when compared to coating directly at high antibody concentrations. However, while the maximal amount of active antibody on the surface can only be increased modestly, we observed that with directed immobilization, much lower antibody concentrations are required for similar binding capacity. For example, 5 μg/mL of SpyTagged Fab coupled for one hour to SpyCatcher3 coated at 5 μg/mL results in a similar amount of active antibody on the surface as 40 μg/mL of Fab coated overnight directly to the plate. Interestingly, we also observed that the binding curve for the direct coating to the plates is much steeper than the curves for SpyTag/SpyCatcher mediated immobilization of antibody. This suggests that the amount of active antibody on the surface can be much better controlled through use of directed immobilization than through direct coating, as variability in coating concentration has less impact on the amount of coated antibody.

**Figure 4:**
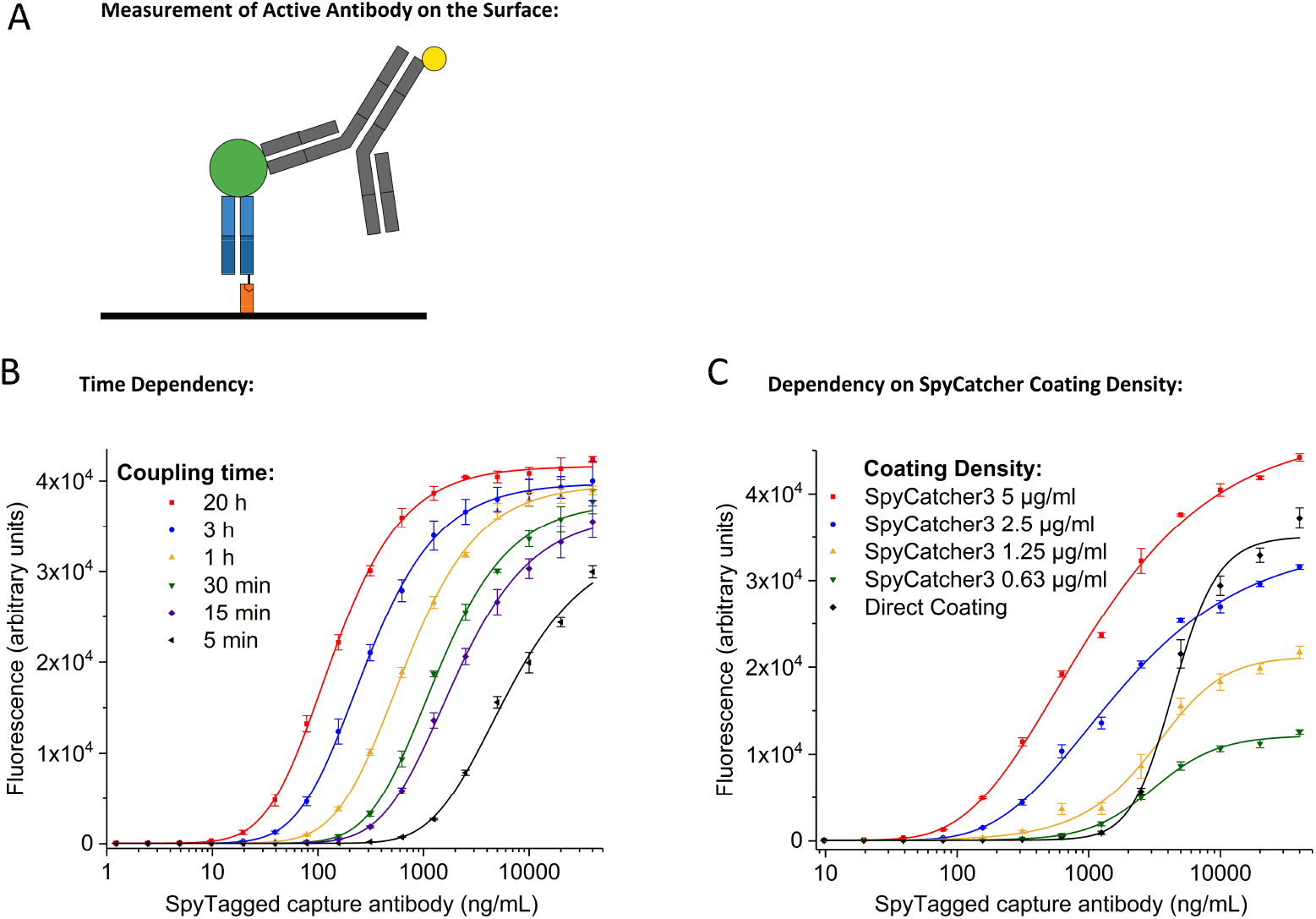
Surface Immobilization via SpyTag/SpyCatcher. **A**: Scheme of sandwich ELISA assay to measure antigen binding capacity of immobilized antibody. SpyCatcher3 (orange) is immobilized to the surface and SpyTagged Fab (blue, format Fab-SpyTag2-His) is immobilized by coupling. FLAG-tagged antigen (green) is detected via an HRP-labeled anti-FLAG IgG (grey). **B**: Time dependency of coupling of SpyTagged Fab to a surface coated with SpyCatcher3 (coating concentration: 5 µg/mL), measured by its capability of binding antigen (adalimumab in Fab-FLAG-His format, 5 µg/mL). **C**: Measurement of active antibody on the surface as achieved by direct coating of the Fab (black curve) vs. coupling to SpyCatcher3 coated at varying densities (colored curves). Binding capacity was measured as before. Error bars are standard deviations from three replicates.

Next, we tested the usefulness of Fab-SpyTag-SpyCatcher coupled products in flow cytometry. For this purpose, we cloned and expressed a mouse antibody against human CD45 (clone F10-89-4^29^) in the Fab-FLAG-SpyTag2-His format and coupled these antibodies with BiCatchers conjugated to phycoerythrin (PE). When we used these coupled products with Jurkat cells for direct detection in flow cytometry, we obtained stronger signals than with commercially available PE-labeled IgGs of the same clone (Figure 5A).

**Figure 5:**
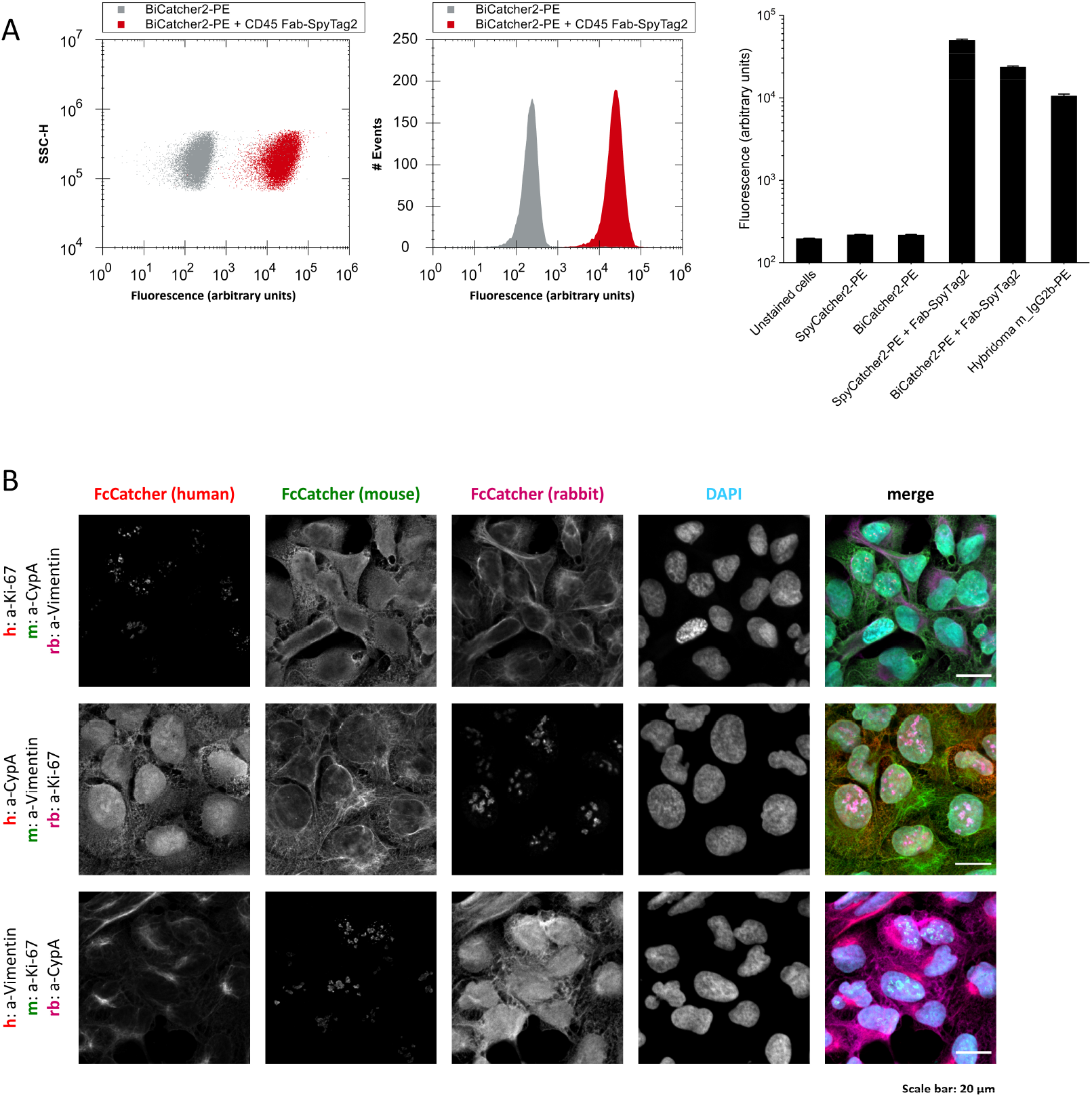
Cell-based Assays with Assembled Modular Antibodies. **A**: Flow cytometry of Jurkat cells with an anti-CD45 mouse Fab-FLAG-SpyTag2-His coupled to BiCatcher2-PE in comparison with BiCatcher2-PE alone. Left: scatter plot of fluorescence intensity (488 nm excitation, 585 nm emission) vs. side scatter height comparing BiCatcher-PE with and without coupled anti-CD45 Fab. Middle: Histogram of the events from the scatter plot. Right: Mean intensities obtained from the Fab coupled to SpyCatcher2-PE or BiCatcher2-PE in comparison with PE-labeled mouse-hybridoma derived IgG and background controls. 8 replicates per measurement were performed, error bars are standard deviations from these 8 independent measurements. **B:** Immunofluorescence confocal microscopy demonstrating complete permutability of antibody specificity and Fc species origin used for detection. Antibodies in Fab-FLAG-SpyTag1-His format against Ki-67, cyclophilin A, and vimentin were coupled with FcCatchers with human, mouse, and rabbit Fc sequences in all possible combinations and incubated with fixed U2OS cells. Detection was performed with secondary antibodies raised against human, mouse, or rabbit Fc domains, and fluorescently labeled with Alexa Fluor 594 (AF594), DyLight488, or AF647, respectively. Individual fluorescence channels as well as a DAPI channel are shown, together with a merged image. Scale bars represent 20 µm.

The superior performance of both PE-labeled and HRP-labeled SpyCatcher constructs compared to directly labeled antibodies is probably due to the fact that labeling of the SpyCatcher molecules can be optimized to achieve a consistent degree of labeling, and there is no risk of labeling at the antigen binding site, as the antibody moiety is coupled via the SpyTag afterwards.

FcCatchers with Fc domains from different species should allow multiplex immunofluorescence for simultaneous detection of multiple target proteins through use of commonly available anti-Fc secondary reagents. Indeed, using human antibodies in the Fab-SpyTag format against cyclophilin A, vimentin, and Ki-67 and FcCatchers with the Fc domains of human, mouse, and rabbit IgG allowed simultaneous visualization of the subcellular location of these proteins in U2OS cells with species-specific secondary detection antibodies (Figure 5B). Immunofluorescence staining worked in all permutations of antibodies and Fc species, underlining the flexibility of the SpyCatcher-based multiplex assays.

### Dual-Format High-Throughput Screening

The modular antibody concept is not restricted to purified antibodies but can also be applied in high throughput to antibodies in lysates. We developed a protocol to change the oligomeric state of the subcloned, uncharacterized antibodies from the HuCAL PLATINUM® phage display library^30^ in *E. coli* lysate without purification. This enables us to screen the relative binding strength of antibodies in 384 microwell plates in both monovalent and bivalent form at the same time, allowing us to select optimal antibodies for a given application. We chose BiCatcher3 (BiCatcher based on SpyCatcher3) for fast coupling at low concentrations. As the concentration of antibodies is unknown in this setting, we first optimized the BiCatcher3 concentration added to the antibody-containing *E. coli* lysates in a dilution series of known GFP binders in the Fab-FLAG-SpyTag2-His format and assayed the coupled antibodies by ELISA (Figure 6A). The antibodies chosen were of very low affinity to maximally benefit from avidity effects. The signal intensity reaches a maximum with increasing BiCatcher concentrations, corresponding to the formation of bivalent (Fab)_2_-BiCatcher before it declines again, when not enough free Fab is available to react with the second SpyCatcher domain of BiCatcher. We chose 60 nM BiCatcher3 as the optimum at which the concentrations of most Fabs and BiCatcher should be closely matched. To demonstrate parallel high-throughput screening of binders in monovalent and bivalent format, we subcloned the output of a low stringency phage display selection against GFP into the Fab-FLAG-SpyTag2-His format. Such a selection can be useful for specific purposes^31^ and was chosen here to model a difficult selection with a limited numbers of screening hits. Expression and lysis of randomly picked clones and controls was performed in 384 microwell plates and a part of the lysate was incubated with BiCatcher3. Both monovalent Fab containing lysate and lysate with coupled bivalent Fab were then screened in parallel by ELISA (Figure 6B). The BiCatcher3 treated lysates showed much higher signals throughout and a striking increase in the number of hits, demonstrating that SpyTag technology could indeed be used to screen monovalent and bivalent binding in parallel, in high throughput, on the same set of unsequenced, unpurified antibodies.

**Figure 6:**
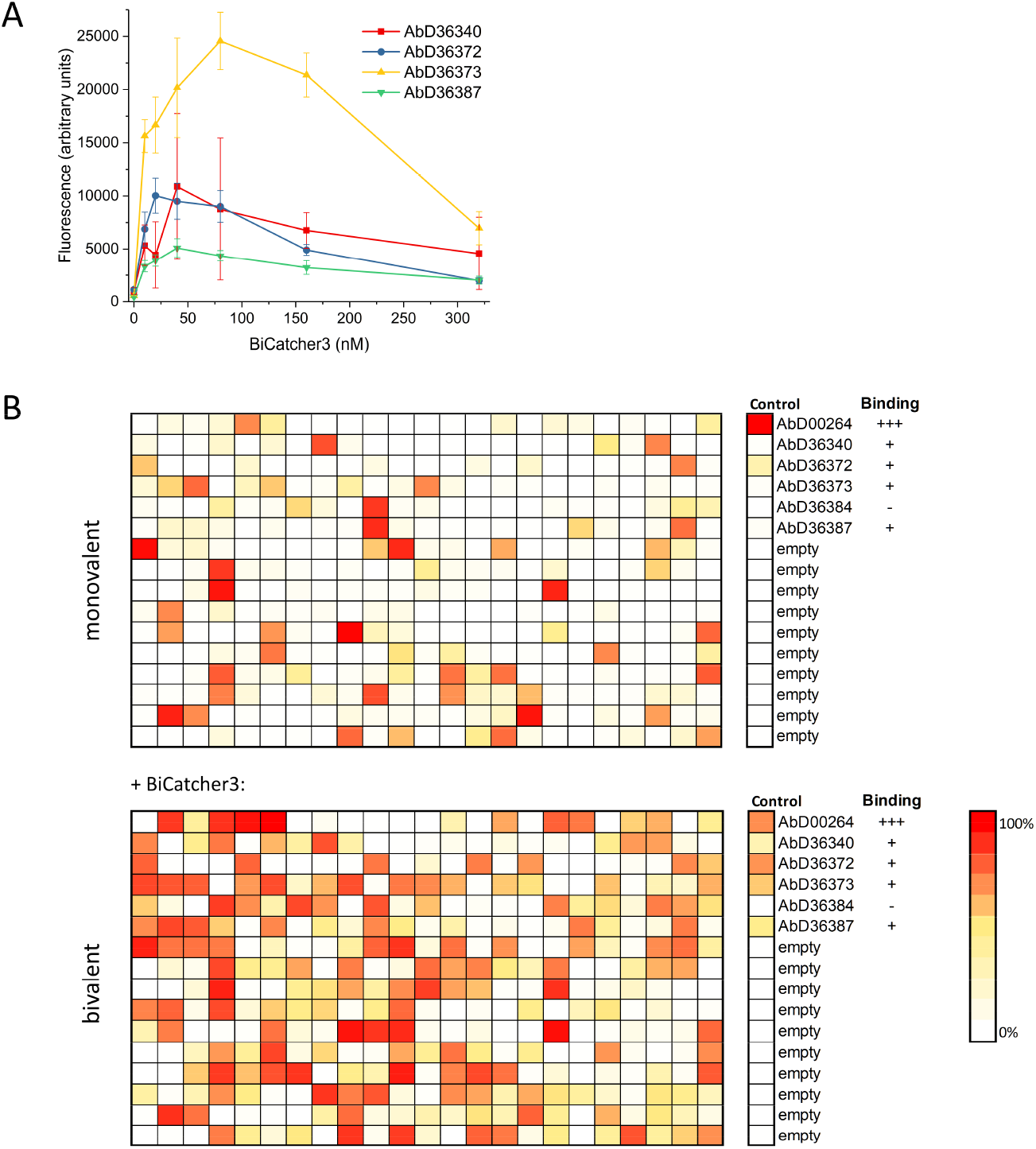
High-Throughput Screening of Antibodies in Both Monovalent and Bivalent Format. **A:** Establishing the optimal concentration for coupling of Fabs in crude bacterial lysate with BiCatcher3. Crude lysate of microscale (80 µL) expression cultures from four low-affinity GFP antibodies (clones AbD36340, AbD36372, AbD36373, AbD36387) was incubated with increasing concentrations of BiCatcher3 for 90 minutes and screened by ELISA with GFP as antigen. Error bars are standard errors of the mean from 8 replicates per data point. **B:** Screening of 368 uncharacterized anti-GFP antibodies (Fab-FLAG-SpyTag2-His format) in crude bacterial lysate by ELISA against GFP. Top: ELISA results from screening a 384 microwell plate from crude bacterial lysate without further treatment. Bottom: ELISA results from screening of the identical lysates, preincubated with 60 nM BiCatcher3 for 1.5 hours. The last row contains controls as indicated, including the binders from subfigure A, as well as empty wells to determine the background signal. Screening hits are color coded according to relative ELISA signal strength according to the heat map.

## Discussion

The SpyTag technology has been successfully adopted for a broad range of applications due to its simple, fast, robust, and quantitative protein ligation reaction^14, 15^. In this study, we show that all published variants of the SpyTag are prone to cleavage by proteases when expressed in the periplasm of *E. coli*.

We offer a solution to this problem by identifying and removing the responsible proteases from *E. coli*, or alternatively, in the case of the initial version SpyTag1, by protecting the SpyTag from physical access by the protease. Expression in knockout *E. coli* cells is robust, delivers high yields, and we expect widespread adoption of this method.

Periplasmic protein expression occurs under oxidizing conditions and is essential for bacterial expression of most proteins containing disulfide bonds, among which antibodies are an important group of considerable academic and commercial interest. With this work, we open up the SpyTag technology for this class of bacterially produced proteins. We have used the methods described here to produce over 1,000 SpyTagged human Fab antibodies as well as mouse Fabs, scFvs, and MBP, making us confident in the robustness and universality of these approaches.

The proteolytic susceptibility of the SpyTag in the periplasm has not been described in the literature. SpyTagged antibody fragments have been expressed in *E. coli* avoiding functional periplasmic expression, such as refolding of insoluble cytoplasmic inclusions^32^ or use of antibody-SpyCatcher fusion proteins^33, 34^. Alternatively, other hosts such as yeast^35^ or mammalian cells^34, 36^ have been used. Nevertheless, there is a small number of studies in which periplasmic expression of SpyTagged proteins were described^16, 17, 18^. Interestingly, during our initial investigation, we also discovered one antibody which could be produced in our standard *E. coli* expression strain, albeit with low yields. Furthermore, we show that cleavage is not completely quantitative with the expression conditions we used, as a small fraction of Fab with full-length SpyTag remains (Supplementary figure S6). We therefore suspect that the efficiency of SpyTag cleavage is to some extent antibody sequence-dependent. However, in our experience, the vast majority of SpyTagged antibody fragments are susceptible to proteolytic cleavage by periplasmic proteases and require mitigation strategies for bacterial expression at useful concentrations.

Modularity is an important concept in the reductionist understanding of biology^37^ as well as in the design of technology, allowing for an efficient reuse of parts in different contexts^38^. In this study, we describe the construction of a highly modular system for antibody reagents based on the SpyTag technology. We have generated a large number of standardized parts that can be used to convert SpyTagged antibody fragments into diverse oligomeric states, equip them with Fc regions of different species/isotypes, or site-specifically label them with probes or enzymes – within minutes to an hour. We show that these assembled modules behave similarly in standard assays to the equivalent antibodies generated by conventional means. Thus the time and effort to convert an antibody to a new format for a custom application is reduced by orders of magnitudes, from weeks to minutes. In this way, the SpyTag technology allows rapid prototyping of antibodies and antibody fusion proteins for a wide variety of applications. Furthermore, we show benefits of using SpyTagged antibodies in assay design, where the oriented immobilization of capture antibodies simultaneously increases assay robustness and reduces the amount of antibody required for ELISA. Other groups have reported the increase of assay sensitivity using such approaches^17^, whereas we see the strengths in this strategy additionally in convenience, limited assay costs, and increased reproducibility.

In contrast to most chemical conjugations, the SpyTag-SpyCatcher coupling reaction is site-specific, resulting in a defined antibody product with high batch-to-batch consistency. Each antibody carrying a SpyTag can be coupled to each one of a large portfolio of pre-built labeled or modified SpyCatchers leading to a multitude of experimental options. We believe that these advantages will stimulate others to clone antibodies from hybridomas in order to make the reagent amenable to the SpyTag technology. Thus, this platform is also a contribution to the ongoing call for improvement of antibody quality by switching to recombinant antibody expression, and might replace poorly characterized antibodies with better defined recombinant versions^39, 40^.

While there are multiple efforts towards using SpyTag technology for vaccine production^41, 42^, the modular antibodies presented in this study will not be suitable as therapeutic antibodies due to the expected strong immune response against SpyCatcher, which is essentially a domain from a *Streptococcus* surface protein. To develop modular therapeutic antibodies, a non-immunogenic protein ligation module would have to be developed.

Finally, we have demonstrated additional advantages of using SpyTagged antibody fragments during antibody discovery. For *in vitro* selection systems, particularly phage display^43^, we show that SpyTag technology offers the possibility to modify the oligomeric state of uncharacterized antibodies in crude bacterial lysates after selection by coupling with oligomeric SpyCatchers. Being able to effortlessly change the valency of uncharacterized antibodies after expression offers a new degree of freedom for the screening of selection outcomes, allowing for bivalent cross-reactivity profiling while still maintaining monovalency for affinity determination. Adding an FcCatcher to the lysates will similarly allow high-throughput functional screening in assays that require the Fc moiety such as antibody-dependent cellular cytotoxicity (ADCC) assays^44^. Furthermore, the addition of an antibody-SpyCatcher fusion protein during the selection step could be used to select for bispecific antibodies, similarly to a recently developed intein-based approach^45^.

In conclusion, the use of SpyTag technology enables rapid and economic development of diagnostic and research antibodies with unprecedented flexibility in tags and labels. Moreover, its potential for rapid prototyping will accelerate preclinical drug discovery research.

## Methods

### Expression Plasmids

All bacterial SpyCatcher expression plasmids used the pET28a vector (Merck KGaA) as backbone. For monomeric SpyCatcher expression plasmids, the coding sequences of SpyCatcher1^11^, SpyCatcher2^12^, or SpyCatcher3^13^ was cloned in frame with coding sequences for an N-terminal hexahistidine tag and a linker (DYDIPTTENLYFQG). BiCatcher expression plasmids carried a sequence coding for the same N-terminal sequence followed by the coding sequence for a second C-terminally attached SpyCatcher in frame, connected by a linker sequence coding for a glycine-serine linker (GSSGSGGGGSGGGGSG). Expression plasmids for MultiCatchers were constructed with identical linkers, however sequences coding for a hexahistidine tag followed by a Strep tag II were attached at the 3’ end of the open reading frame. Mono-SpyCatcher variants carrying an additional cysteine were constructed by inserting the coding sequence for a cysteine encompassed by two glycines directly at the 5’ end of the SpyCatcher coding sequence. BiCatcher carrying three cysteines was generated by adding an N-terminal cysteine as above, replacing the codon for the central serine in the linker with a cysteine codon, and adding a C-terminal cysteine by placing a sequence coding for GGC at the very 3’ end of the open reading frame.

For expression in cell culture, FcCatcher proteins were cloned into the pMax (MorphoSys) vector. The sequence coding for SpyCatcher3 followed by the sequence coding for a GSSGS linker was inserted 5’ to the sequence coding for the antibody hinge region followed the sequence coding for the Fc part of the various isotypes and species.

For bacterial antibody fragment expression, the pMORPHx11 (MorphoSys) or pBBx1/pBBx2 (Bio-Rad) plasmids were used. The periplasmic export signal sequences and C-terminal tag sequences are provided in the supplement. The sequences of the MBP proteins are listed in the supplement as well. pMorphx11 was used for periplasmic expression, pET28a for cytosolic expression.

For each construct, codon-optimized double-stranded gene fragments with appropriate restriction sites were ordered from Twist Biosciences or Eurofins Genomics and cloned into the appropriate plasmids.

Full-length immunoglobulins were expressed using the pMorph2 or pMorph4 plasmids (MorphoSys).

### Protein Expression and Purification

Human Fab fragments selected from the HuCAL GOLD^46^ or PLATINUM library^30^ were expressed and purified by Ni-NTA affinity chromatography as described^23^. Briefly, bacteria (*E. coli* TG1 F- or mutant strains SK4/SK13) were grown in 2xYT at 30°C until an OD600 of about 0.5 was reached. Antibody expression was induced with 1 mM IPTG, followed by overnight incubation at 30°C (27.5°C for SK13) at 250 rpm. Cells were harvested by centrifugation, lysed with BugBuster (Merck KGaA) supplemented with 2 mg/mL lysozyme (Merck KGaA), 20 units/mL Benzonase (Merck KGaA), and protease inhibitors (Roche Complete EDTA free), and passed over Ni-NTA agarose (Qiagen). Proteins were eluted with buffer containing 250 mM imidazole, 500 mM NaCl, and 20 mM NaH_2_PO_4_ pH7.4. Buffer was exchanged to PBS with PD10 (GE Healthcare) or CentriPure P96 (emp Biotech) columns. Murine Fab fragments were expressed as above. Following Ni-NTA purification, murine Fab fragments were buffer exchanged into MES 20 mM, NaCl 20 mM, pH 6.27 and applied on a Bio-Scale Mini UNOsphere S (Bio-Rad). Elution was accomplished with a gradient to 50% of MES, 20 mM, NaCl 1 M, pH 6.27. SpyCatcher and BiCatcher proteins were expressed in *E. coli* BL21 cells and purified by Ni-NTA affinity chromatography as described above, however instead of BugBuster, IMAC-RB (20 mM NaH_2_PO_4_, 500 mM NaCl, 10 mM imidazole, supplemented with lysozyme, Benzonase and protease inhibitors as above) was used as lysis buffer. BiCatcher2 was expressed overnight at 20°C to minimize cleavage of the glycine-rich linker. For analysis of expression via immunoblotting or ELISA, samples in lysis buffer without further purification were used. FcCatchers and full-length immunoglobulins were expressed and purified as described^47^. Briefly, HKB11 cells^48^ were transiently transfected and grown in shake flasks. Proteins were purified from supernatants by protein A affinity chromatography using ÄKTA (GE Healthcare) or NGC (Bio-Rad) chromatography systems and rebuffered to PBS.

### Protein Labeling

Labeling of proteins with engineered cysteines was performed by standard maleimide chemistry^49^. Briefly, proteins were freshly reduced with 1 mM TCEP or 1 mM DTT in PBS for 60 minutes at room temperature. The reducing agents were removed by rebuffering into PBS with desalting columns (GE Healthcare PD10) and freshly reduced proteins were mixed with malemide-labeled PE (Expedeon) or HRP (Thermo-Fisher Scientific) according to the manufacturer’s instructions. The mixture was incubated for one hour at room temperature or overnight at 4°C and quenched with 1 mM cysteine. Alternatively, we used commercially available BiCatcher-HRP or BiCatcher-PE (Bio-Rad).

### Protein Ligation (SpyTag-SpyCatcher Coupling Reaction)

SpyTagged proteins and SpyCatcher were mixed in PBS and incubated usually for one hour at room temperature. For mono-SpyCatcher proteins, a molar excess of 25% of SpyCatcher over SpyTag was chosen to ensure complete coupling of SpyTagged proteins. For multivalent SpyCatcher proteins, a 25% molar excess of SpyTagged proteins over SpyCatcher sites was chosen to ensure complete multivalency of all SpyTag-SpyCatcher coupled products. Typically, couplings were performed at monovalent SpyCatcher concentrations ranging between 1 and 10 µM. For some experiments, excess free SpyTagged proteins were removed by passing the reaction mixture over a column (Profinity resin, Bio-Rad) on which SpyCatcher3 was immobilized via expoxide chemistry according to the manufacturer’s instructions.

### ELISA

ELISAs were performed as described^47^. Briefly, antigens or capture antibodies were coated in PBS overnight onto 384 microwell plates (Maxisorp, Thermo Fisher Scientific), washed 5 times with PBST (PBS, 0.05% Tween20), and blocked for one hour with PBST/5% BSA, PBST/5% milk powder or ChemiBLOCKER (Merck KGaA). After further washing, analytes were titrated in a dilution series in an appropriate buffer (e.g. PBST/1%BSA) and incubated for one hour. Where necessary, detection antibodies were bound consecutively in the same manner. Detection was performed via QuantaBlu fluorogenic peroxidase substrate (Thermo Fisher Scientific). For absolute Fab concentration determination by ELISA (Figure 1E), a Fab standard of known concentration was measured in a dilution series in 4 replicates and the linear region was used to fit a linear equation. 3-4 values from within this linear range were selected to measure the concentration of unknown Fabs (in duplicates), and an average of these values was calculated, yielding the absolute Fab concentration. Detection antibodies used: anti-human F(ab’)_2_ IgG-HRP (STAR126P, Bio-Rad), anti-His5-Tag IgG-HRP (MCA5995P, Bio-Rad), anti-FLAG-Tag (M2) IgG-HRP (A8592, Sigma).

### Immunoblotting

SDS PAGE precast gels (TGX, Bio-Rad) were blotted via semi-dry transfer onto nitrocellulose or PVDF membranes using Trans-Blot Turbo Transfer packs (Bio-Rad) and the Trans-Blot Turbo Transfer system (Bio-Rad) with default settings. Membranes were blocked for one hour in TBST (TBS, 0.05% Tween20) containing 5% milk powder, washed 3 times with TBST, and incubated with primary antibody for one hour. If necessary, the blots were again washed 3 times with TBST and incubated with a secondary HRP-labeled antibody for one hour. Blots were washed 3 times with TBST and then incubated with Clarity ECL substrate (Bio-Rad). Blots were imaged using a ChemiDoc MP gel doc system (Bio-Rad). Detection antibodies used: anti-human F(ab’)_2_ IgG-HRP (STAR126P, Bio-Rad), anti-His5-Tag IgG-HRP (MCA5995P, Bio-Rad), anti-FLAG-Tag (M2) IgG-HRP (A8592, Sigma).

### Mass Spectrometry

Mass spectrometry was performed by TOPLAB GmbH, Martinsried, Germany. Briefly, for mass determination, Ni-NTA purified proteins were desalted and co-crystalized with sinapinic acid matrix. Mass spectroscopy was performed on a 4800 MALDI TOF/TOF Analyzer (AB Sciex) in linear mode from 5000 – 50 000 m/z with an estimated accuracy of 1000 ppm. The instrument was calibrated with a protein standard (Bruker) containing cytochrome C and apomyoglobin. Samples were analyzed with Data Explorer (Applied Biosystems). For tryptic digests, samples were reduced in 6 M urea, 0.3 M ammonium bicarbonate, 4 mM DTT, pH 8.5 for 30 minutes at 55°C and alkylated with 8 mM iodoacetamide for 15 minutes at 25°C. Samples were then diluted to 2 M urea and digested overnight with trypsin. Samples were desalted and co-crystalized with alpha-cyano-4-hydroxycinnamic acid matrix. The instrument was run in reflection mode from 700 – 7000 m/z. Peptide masses were compared to expected masses with Mascot (Matrix Science).

### Screening

To determine the optimal concentration of BiCatcher3 for screening, four low affinity anti-GFP Fabs (AbD36340, AbD36372, AbD36373, AbD36387) and one non-binding Fab (AbD36384) were subcloned into the Fab-FLAG-SpyTag2-His format and transformed into SK13 cells. For each Fab, 64 wells of a 384 microwell plate containing 80 µL of 2xYT medium supplemented with 34 µg/mL chloramphenicol, 1% glucose, 10% glycerol were inoculated with single colonies and incubated for 5 hours at 37°C shaking at 400 rpm. 5 µL of these cultures were used to inoculate a new 384 microwell plate with each well containing 55 µl of 2xYT supplemented with 34 µg/mL chloramphenicol, 0.1% glucose, 0.8 mM IPTG. This new plate was then incubated shaking at 400 rpm for 2 hours at 37°C followed by overnight incubation at 22°C. The next day, cells were lysed by adding to each well 15 µL lysis buffer (0.4 M borate pH 8.0, 320 mM NaCl, 4 mM EDTA, 2.5 mg/mL lysozyme, 12.5 units/mL Benzonase) and incubating for 1 hour at 22°C at 400 rpm. For coupling, 20 µL of lysate from each well was transferred onto a 384 microwell plate containing 20 µL of BiCatcher3 in PBS in a dilution series from 640 nM to 10 nM or PBS alone and incubated for 1.5 hours at room temperature, with 8 replicates per antibody and dilution step. Coupled lysates were then transferred onto a plate coated with GFP at 5 µg/mL and blocked with PBST/5% milk powder. ELISA was performed as described above, with goat anti-human IgG F(ab’)_2_-HRP (STAR126P, Bio-Rad) as the detection antibody.

For screening, the output of the third round of a phage display selection against GFP performed with the HuCAL PLATINUM library was subcloned from the display vector into the pBBx1 expression vector in the FLAG-SpyTag2-His format and transformed into SK13 cells. 368 clones were picked and used to inoculate a 384 microwell plate as described above. As controls, additional wells were inoculated with SK13 cells expressing a high-affinity anti-GFP antibody (AbD00264) or the five antibodies from the coupling optimization. Growth and lysis were carried out as described above. Coupling was performed as described above, albeit with a constant final concentration of 60 nM BiCatcher3 in the coupling reaction. For ELISA, the number of wash repeats was doubled in each step (i.e. 10x washing instead of 5x), and otherwise performed as described above.

### *E. coli* Knockout

Construction of knockout strains was performed as described^21^. Briefly, the PCR cassette tsp_kanR_pKD13 for deletion of the tsp-locus was amplified by using pKD13 as template and the primers 119_SKE and 120_SKE. TG1F-harboring pKD46 were made electrocompetent (expression of recombinase was induced by addition of 50 mM arabinose) and transformed with the PCR cassette tsp_kanR_pKD13. After integration and excision of the kanR-cassette by transformation with pCP20, Δtsp-mutants were identified by colony PCR using the primers 88_SKE and 89_SKE. The deletion scar was verified by sequencing and the final strain was termed SK4. For deletion of the ompT-locus, the PCR cassette ompT_kanR_pKD13 was amplified with the primers 127_SKE, 128_SKE, and pKD13 as template. SK4 harboring pKD46 were made electrocompetent and transformed with the PCR cassette ompT_kanR_pKD13. ΔompT-mutants were identified by colony PCR using the primers 135_SKE and 136_SKE. Furthermore, the Δtsp-locus was analyzed a second time for correctness of deletion. The deletion scar ΔompT was verified by sequencing and the final strain with Δtsp and ΔompT-locus was termed SK13. The primer sequences used are listed in Supplementary table 1.

### Cell Culture

For immunofluorescence (IF) experiments, U2OS cells (human osteosarcoma, ATCC-HTB-96) were grown in DMEM medium containing 4.5 g/L glucose, 1 mM pyruvate, and GlutaMAX (Gibco) supplemented with 10% FBS (ultra-low IgG, Gibco). Cells were grown in T75 flasks (Sarstedt) at 37°C in humidified atmosphere containing 5% CO_2_ and routinely passaged by trypsinization with 0.25% trypsin-EDTA (Gibco).

For flow cytometry experiments, Jurkat cells were grown in ATCC-modified RMPI 1640 medium (Gibco) supplemented with 10% FBS (ultra-low IgG, Gibco). Cells were routinely grown in 125 mL flasks (Corning), shaken at 100 rpm on an orbital shaker with 25 mm stroke at 37°C in humidified atmosphere containing 5% CO_2_, and routinely passaged to maintain cell density below 2.5 x 10^6^ cells/mL.

### Immunofluorescence

3.75 x 10^4^ U2OS cells per well were seeded in 12-well chamber slides with removable silicone gaskets (Ibidi). After 24 hours growth, cells were washed twice with pre-warmed PBS (containing calcium and magnesium salts for this washing step), fixed 15 min in 4% PFA/PBS, washed twice with PBS, incubated with ice-cold methanol for 10 min at -20°C, washed 3 times with PBS, treated 5 min with 0.2% Triton X-100 in PBS, washed once with PBS, and incubated 36 hours at 4°C in blocking solution (5% BSA, 0.1% Tween20, PBS). After blocking, slides were equilibrated to room temperature and cells were treated with mixtures of primary reagents in blocking solution. Applied primary reagents were human Fab-FLAG-SpyTag-His against human Ki-67, human cyclophilin A and human vimentin, coupled to either human IgG1-, mouse IgG2a-, or rabbit IgG-FcCatchers. The concentration of coupled products for anti-Ki-67 and anti-vimentin was 33 nM, whereas 167 nM coupled FcCatchers were used for anti-cyclophilin A. After 3 hours incubation at room temperature, cells were washed 4 times with wash buffer (0.1% Tween20 in PBS) and incubated for 1 hour at room temperature with a mixture of detection reagents in blocking buffer. Detection reagents consisted of (i) goat anti-human IgG (Fcγ specific) F(ab’)_2_ AF594 conjugate (Jackson ImmunoResearch 109-586-170), (ii) goat anti-mouse IgG (H+L), DyLight488 conjugate (Bio-Rad STAR117D488GA), (iii) donkey anti-rabbit IgG (H+L), AF647 conjugate (Jackson ImmunoResearch 711-605-152) and (iv) DAPI, 333 ng/mL (Merck KGaA). After incubation, cells were washed 4 times with wash buffer, once with PBS, and mounted in ProLong Gold Antifade Mountant (Molecular Probes), cured overnight at room temperature, and stored at 4°C.

Cells were imaged on Zeiss LSM 880 confocal microscope with an EC Plan-Neofluar 40x/1.30 immersion lens. 16 bit mode, 1024×1024 frame size and 4 line averages were used. Pinhole size, laser power, and detector gain were separately adjusted for each of the channels. Brightness and contrast were adjusted with ImageJ (NIH) and representative regions of interest were selected.

### Flow Cytometry

Cultivated Jurkat cells were collected and washed 3 times in flow buffer (PBS without Ca/Mg, 3% FBS, Gibco). Cell concentration was measured, diluted to 1.5 x 10^6^ cells/mL, and 30,000 cells per well were seeded into a V-shaped 384 microwell plate (Greiner). Respective PE-Catcher-coupled antibodies were added at a final concentration of 10 nM and cells were incubated for one hour in the dark, followed by 3 additional washing steps. All cell handling was performed at 4°C or on ice whenever possible. After adjusting the volume to 40 µL per well with flow buffer, approximately 2000 cells per well were measured with the IntelliCyt® iQue Screener flow cytometer (Sartorius) and analyzed using ForeCyt analysis software (Sartorius). Results shown are the accumulation of 8 replicates per condition.

### Nanoscale Differential Scanning Fluorimetry (nanoDSF)

Samples were diluted in PBS, loaded into sample capillaries by capillary tension, and measured in duplicates in a Prometheus NT.48 (NanoTemper) with the following settings: 35-40% excitation power, temperature slope 0.5-0.7°C/min, starting temperature 20°C, end temperature 95°C.

## Acknowledgements

We thank our colleagues in the protein production teams, especially Waldemar Preis, for the expression and purification of the proteins used in this study. We thank Lena Schmohl for her help during the initial phase of this project. We thank Farida Hellal for help with microscopy. We thank Antje Hentrich for illustrations. We thank Amanda Turner and Philipp Diebolder for careful reading of the manuscript and Christian Frisch for helpful discussions. Mass spectrometry was carried out by TOPLAB GmbH, Martinsried, Germany.

## Contributions

C.H., S.-J.K., M.P., M.C., and H.H. performed experiments. C.H., S.-J.K., M.P., and F.Y. designed experiments and analyzed the data. A.K. and F.Y. conceived the project. C.H., A.K., and F.Y. wrote the manuscript with input from M.P. and S.-J.K..

## References

1. Borrebaeck CAK. Antibodies in diagnostics – from immunoassays to protein chips. Immunology Today 21, 379–382 (2000).

2. Goldman R. Antibodies: indispensable tools for biomedical research. Trends in Biochemical Sciences 25, 593–595 (2000).

3. Bates A, Power CA. David vs. Goliath: The Structure, Function, and Clinical Prospects of Antibody Fragments. Antibodies (Basel, Switzerland) 8, (2019).

4. Greenfield EA. in Antibodies: A laboratory manual Greenfield EA (Ed.), Ch. 7, 201–301 (Cold Spring Harbor Laboratory Press, 2014).

5. Skerra A, Pluckthun A. Assembly of a functional immunoglobulin Fv fragment in Escherichia coli. Science 240, 1038–1041 (1988).

6. Pluckthun A. Antibodies from Escherichia coli. Nature 347, 497–498 (1990).

7. Manning CF, Bundros AM, Trimmer JS. Benefits and pitfalls of secondary antibodies: why choosing the right secondary is of primary importance. PLoS One 7, e38313 (2012).

8. Hermanson GT. in Bioconjugate techniques Ch. 10, 456–493 (Academic Press, 1996).

9. Adumeau P, Sharma SK, Brent C, Zeglis BM. Site-Specifically Labeled Immunoconjugates for Molecular Imaging—Part 1: Cysteine Residues and Glycans. Molecular Imaging and Biology 18, 1–17 (2016).

10. van Vught R, Pieters RJ, Breukink E. Site-specific functionalization of proteins and their applications to therapeutic antibodies. Comput Struct Biotechnol J 9, e201402001–e201402001 (2014).

11. Zakeri B, et al. Peptide tag forming a rapid covalent bond to a protein, through engineering a bacterial adhesin. Proc Natl Acad Sci U S A 109, E690–697 (2012).

12. Keeble AH, Banerjee A, Ferla MP, Reddington SC, Anuar I, Howarth M. Evolving Accelerated Amidation by SpyTag/SpyCatcher to Analyze Membrane Dynamics. Angew Chem Int Ed Engl, (2017).

13. Keeble AH, et al. Approaching infinite affinity through engineering of peptide-protein interaction. Proc Natl Acad Sci U S A, (2019).

14. Keeble AH, Howarth M. Power to the Protein: Enhancing and Combining Activities using the Spy Toolbox. Chemical Science, (2020).

15. Reddington SC, Howarth M. Secrets of a covalent interaction for biomaterials and biotechnology: SpyTag and SpyCatcher. Curr Opin Chem Biol 29, 94–99 (2015).

16. Alam MK, Gonzalez C, Hill W, Fonge H, Barreto K, Geyer CR. Synthetic Modular Antibody Construction Using the SpyTag/SpyCatcher Protein Ligase System. Chembiochem, (2017).

17. Anderson GP, et al. Oriented Immobilization of Single-Domain Antibodies Using SpyTag/SpyCatcher Yields Improved Limits of Detection. Analytical chemistry 91, 9424–9429 (2019).

18. Kimura H, Asano R, Tsukamoto N, Tsugawa W, Sode K. Convenient and Universal Fabrication Method for Antibody-Enzyme Complexes as Sensing Elements Using the SpyCatcher/SpyTag System. Anal Chem 90, 14500–14506 (2018).

19. Silber KR, Keiler KC, Sauer RT. Tsp: a tail-specific protease that selectively degrades proteins with nonpolar C termini. Proc Natl Acad Sci U S A 89, 295–299 (1992).

20. Hara H, Yamamoto Y, Higashitani A, Suzuki H, Nishimura Y. Cloning, mapping, and characterization of the Escherichia coli prc gene, which is involved in C-terminal processing of penicillin-binding protein 3. J Bacteriol 173, 4799–4813 (1991).

21. Datsenko KA, Wanner BL. One-step inactivation of chromosomal genes in Escherichia coli K-12 using PCR products. Proc Natl Acad Sci U S A 97, 6640–6645 (2000).

22. McCarter JD, Stephens D, Shoemaker K, Rosenberg S, Kirsch JF, Georgiou G. Substrate specificity of the Escherichia coli outer membrane protease OmpT. J Bacteriol 186, 5919–5925 (2004).

23. Knappik A, Brundiers R. in The Protein Protocols Handbook Walker JM (Ed.), 1929–1943 (Humana Press, 2009).

24. Tan LL, Hoon SS, Wong FT. Kinetic Controlled Tag-Catcher Interactions for Directed Covalent Protein Assembly. PLoS One 11, e0165074 (2016).

25. Bae Y, Jang DG, Eom S, Park TJ, Kang S. HRP-conjugated plug-and-playable IgG-binding nanobodies as secondary antibody mimics in immunoassays. Sensors and Actuators B: Chemical 320, 128312 (2020).

26. Real-Hohn A, Groznica M, Loffler N, Blaas D, Kowalski H. nanoDSF: In vitro Label-Free Method to Monitor Picornavirus Uncoating and Test Compounds Affecting Particle Stability. Front Microbiol 11, 1442 (2020).

27. Lin Z, et al. Spy chemistry-enabled protein directional immobilization and protein purification. Biotechnol Bioeng, (2020).

28. Butler JE, et al. The physical and functional behavior of capture antibodies adsorbed on polystyrene. J Immunol Methods 150, 77–90 (1992).

29. Dalchau R, Kirkley J, Fabre JW. Monoclonal antibody to a human leukocyte-specific membrane glycoprotein probably homologous to the leukocyte-common (L-C) antigen of the rat. Eur J Immunol 10, 737–744 (1980).

30. Prassler J, et al. HuCAL PLATINUM, a synthetic Fab library optimized for sequence diversity and superior performance in mammalian expression systems. J Mol Biol 413, 261–278 (2011).

31. Chatterjee T, et al. Direct kinetic fingerprinting and digital counting of single protein molecules. Proceedings of the National Academy of Sciences 117, 22815–22822 (2020).

32. Yumura K, et al. Use of SpyTag/SpyCatcher to construct bispecific antibodies that target two epitopes of a single antigen. The Journal of Biochemistry 162, 203–210 (2017).

33. Alam MK, Brabant M, Viswas RS, Barreto K, Fonge H, Ronald Geyer C. A novel synthetic trivalent single chain variable fragment (tri-scFv) construction platform based on the SpyTag/SpyCatcher protein ligase system. BMC biotechnology 18, 55 (2018).

34. Alam MK, El-Sayed A, Barreto K, Bernhard W, Fonge H, Geyer CR. Site-Specific Fluorescent Labeling of Antibodies and Diabodies Using SpyTag/SpyCatcher System for In Vivo Optical Imaging. Molecular imaging and biology 21, 54–66 (2019).

35. Wichgers Schreur PJ, et al. Multimeric single-domain antibody complexes protect against bunyavirus infections. Elife 9, (2020).

36. Fierer JO, Veggiani G, Howarth M. SpyLigase peptide-peptide ligation polymerizes affibodies to enhance magnetic cancer cell capture. Proc Natl Acad Sci U S A 111, E1176–1181 (2014).

37. Lorenz DM, Jeng A, Deem MW. The emergence of modularity in biological systems. Phys Life Rev 8, 129–160 (2011).

38. Baldwin CY, Clark KB. in Design rules Volume 1. The Power of Modularity Ch. 3, 63–92 (MIT Press, 2000).

39. Bradbury AR, Pluckthun A. Getting to reproducible antibodies: the rationale for sequenced recombinant characterized reagents. Protein Eng Des Sel 28, 303–305 (2015).

40. Gray AC, Bradbury A, Dubel S, Knappik A, Pluckthun A, Borrebaeck CAK. Reproducibility: bypass animals for antibody production. Nature 581, 262 (2020).

41. Thrane S, et al. Bacterial superglue enables easy development of efficient virus-like particle based vaccines. J Nanobiotechnology 14, 30 (2016).

42. Bruun TUJ, Andersson AC, Draper SJ, Howarth M. Engineering a Rugged Nanoscaffold To Enhance Plug-and-Display Vaccination. ACS Nano 12, 8855–8866 (2018).

43. Hentrich C, Ylera F, Frisch C, Ten Haaf A, Knappik A. in Handbook of Immunoassay Technologies Vashist SK, Luong JHT (Eds.), 47–80 (Academic Press, 2018).

44. Xiao X, et al. A high-throughput platform for population reformatting and mammalian expression of phage display libraries to enable functional screening as full-length IgG. MAbs 9, 996–1006 (2017).

45. Hofmann T, et al. Intein mediated high throughput screening for bispecific antibodies. mAbs 12, 1731938 (2020).

46. Rothe C, et al. The human combinatorial antibody library HuCAL GOLD combines diversification of all six CDRs according to the natural immune system with a novel display method for efficient selection of high-affinity antibodies. J Mol Biol 376, 1182–1200 (2008).

47. Harth S, Ten Haaf A, Loew C, Frisch C, Knappik A. Generation by phage display and characterization of drug-target complex-specific antibodies for pharmacokinetic analysis of biotherapeutics. MAbs 11, 178–190 (2019).

48. Cho MS, Yee H, Chan S. Establishment of a human somatic hybrid cell line for recombinant protein production. J Biomed Sci 9, 631–638 (2002).

49. Hermanson GT. in Bioconjugate techniques Ch. 2, 146–168 (Academic Press, 1996).

50. Li L, Fierer JO, Rapoport TA, Howarth M. Structural analysis and optimization of the covalent association between SpyCatcher and a peptide Tag. J Mol Biol 426, 309–317 (2014).

