## Supplementary Data for "Periplasmic Expression of SpyTagged Antibody Fragments Enables Rapid Modular Antibody Assembly"

### Supplementary Material

- **Supplementary Figures 1 – 9**
- **Supplementary Table 1**
- **Supplementary Text 1: Protein Sequences**

### Supplementary Figures

#### Figure S1

Light Chain:
Sequence coverage: 85%
Matched peptides are shown in bold red.

1 **DIELTQPPSV SVSPGQTASI TCXXXXXXXX XXXWYQQKPG QAPVXXXXXX**

51 **XXXXXGIPER FSGSNSGNTA TLTISGTQAE DEADYYCXXX** XXXXXXVFGG

101 GTK**LTVLGQP KAAPSVTLFP PSSEELQANK ATLVCLISDF YPGAVTVAWK**

151 ADSSPVK**AGV ETTTPSKQSN NKYAASSYLS LTPEQWK**SHR **SYSCQVTHEG**

201 **STVEK**TVAPT EA

Heavy Chain:
Sequence coverage: 73%
Matched peptides are shown in bold red.

1 **EVQLVESGGG LVKPGGSLRL SCAASXXXXX XXXXXWVR**QA PGK**GLEXXXX**

51 **XXXXXXXXXX XXXXXXXXRF TISRDDSKNT LYLQMNSLKT EDTAVYYCAR**

101 **XXXXXXXXXX WGQGTLVTVS SASTK**GPSVF PLAPSSKSTS GGTAALGCLV

151 K**DYFPEPVTV SWNSGALTSG VHTFPAVLQS SGLYSLSSVV TVPSSSLGTQ**

201 **TYICNVNHKP SNTK**VDKKVE PKSEFHHHHH HGAPGAHIVM VDAYKPTK

**Figure S1: Tryptic Digest Consistent with C-terminal Cleavage of Fab-His-SpyTag1**

Peptide mass fingerprint analysis after tryptic digest of a Fab in His-SpyTag1 format, purified from TG1 F-cells. The identical protein batch was used in the full mass determination in Figure 1C. Matched peptides are shown in red. CDR amino acids were replaced by X for this figure. The N-terminal peptide of the heavy chain was found, whereas the C-terminal peptides were not observed, indicating that the truncation of the heavy chain observed in full mass determination (Figure 1C) occurs at the C-terminus.

Figure S2

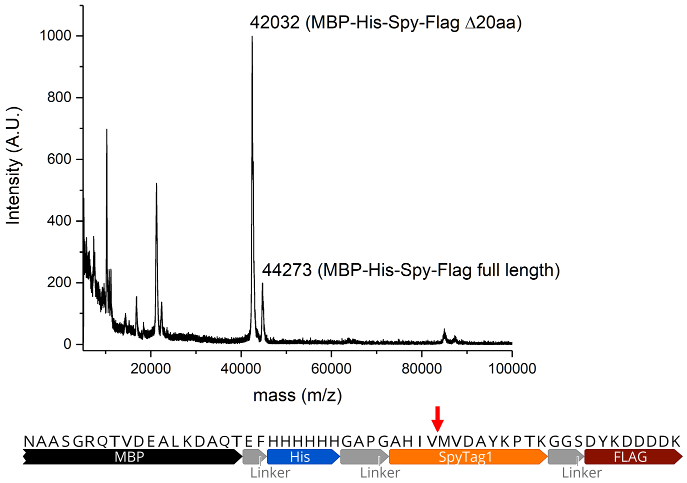


**Figure S2: Mass Spectrometry of Cleaved Periplasmically Expressed MBP-SpyTag1**

Linear mode MALDI mass spectrum (top) of a Ni-NTA affinity-purified MBP-His-SpyTag1-FLAG and C-terminal sequence (bottom). Expected protonated masses: 44,241 Da (full-length), 42,011 Da (20 amino acid C-terminal truncation). The putative cleavage site is indicated by a red arrow.

Figure S3

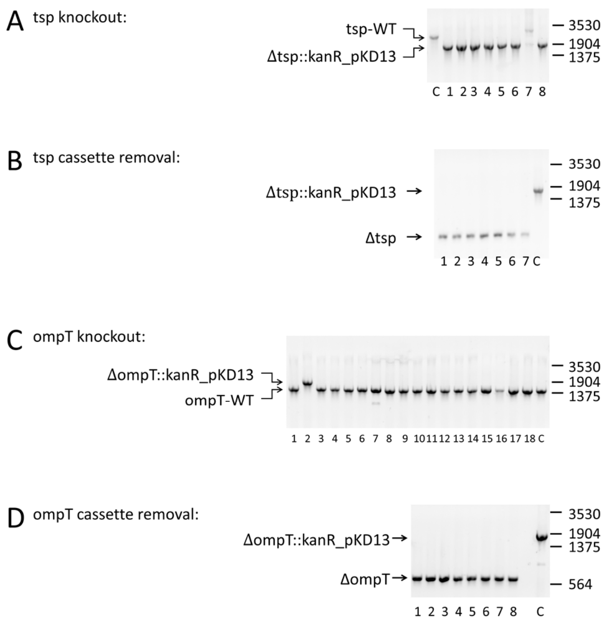


**Figure S3: Generation of *E. coli* Mutant Strains SK4 and SK13.**

**A**: Colony PCR screening of 8 clones showing the replacement of the wildtype tsp locus with a kanamycin resistance cassette in 7/8 clones. Lane C: wild type TG1 F- control. **B**: Colony PCR screening showing the excision of the resistance cassette in the former tsp locus. All 7 clones show proper excision, yielding SK4 (TG1 F- Δtsp). Lane C: Control with the kanamycin resistance cassette in the tsp locus. **C**: Colony PCR screening of 18 clones for the replacement of the wildtype ompT locus with a kanamycin resistance cassette. Only one clone shows insertion of the resistance cassette in the ompT locus. The low efficiency is likely due to the presence of an excision scar sequence at the former tsp locus which is also homologous to the resistance cassette. Lane C: SK4 control. **D**: Colony PCR screening showing the excision of the resistance cassette in the former ompT locus. All 8 clones show proper excision, yielding SK13 (TG1 F- Δtsp ΔompT). Lane C: Control with the kanamycin resistance cassette in the ompT locus. Molecular weights are indicated in units of base pairs.

Figure S4

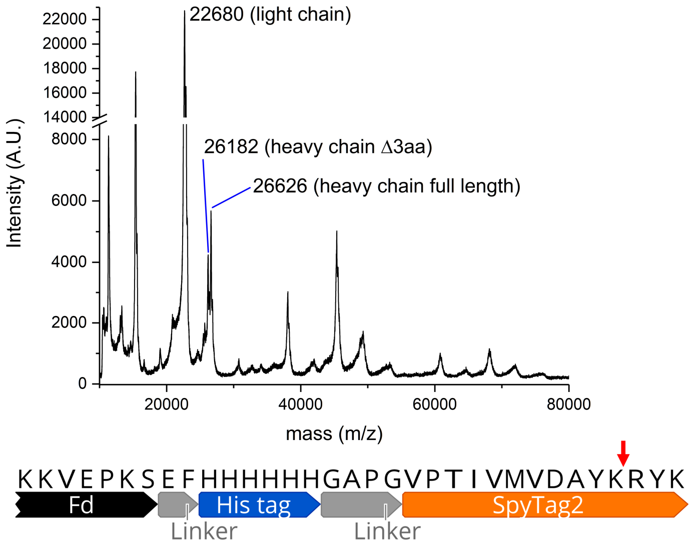


**Figure S4**: **Mass Spectrometry of Cleaved Periplasmically Expressed Fab-His-SpyTag2**

Linear mode MALDI mass spectrum (top) of a Ni-NTA affinity-purified Fab in His-SpyTag2 format (bottom) expressed in SK4. Expected protonated full-length masses: 22,691 Da (light chain), 26,643 Da (heavy chain). Expected protonated mass for the heavy chain with a 3 amino acid C-terminal truncation: 26,195 Da. The relative intensity of peaks does not correspond to abundance of protein species in sample.

Figure S5

A

```
1) MBP Flag-SpyTag-His*: MBP...EALKDAQT - EF - DYKDDDDK - GGS - AHIVMVDAYKPTK - GAP - HHHHHH
2) MBP ST4 SpyTag-His:   MBP...EALKDAQT - AHIVMVDAYKPTK - GAP - HHHHHH
3) ST4:      IGHG1 CH1...KVDKKV - EP - AHIVMVDAYKPTK - GAP - HHHHHH
4) ST4+1:    IGHG1 CH1...KVDKKV - EPK - AHIVMVDAYKPTK - GAP - HHHHHH
5) ST4+2:    IGHG1 CH1...KVDKKV - EPKS - AHIVMVDAYKPTK - GAP - HHHHHH
6) ST4+3:    IGHG1 CH1...KVDKKV - EPKSP - AHIVMVDAYKPTK - GAP - HHHHHH
7) ST4+4:    IGHG1 CH1...KVDKKV - EPKSGG - AHIVMVDAYKPTK - GAP - HHHHHH
8) ST4+5**: IGHG1 CH1...KVDKKV - EPKSEFG - AHIVMVDAYKPTK - GAP - HHHHHH

* produced cytoplasmically
** produced in protease knockout E.Coli SK4
Linker  SpyTag1
```

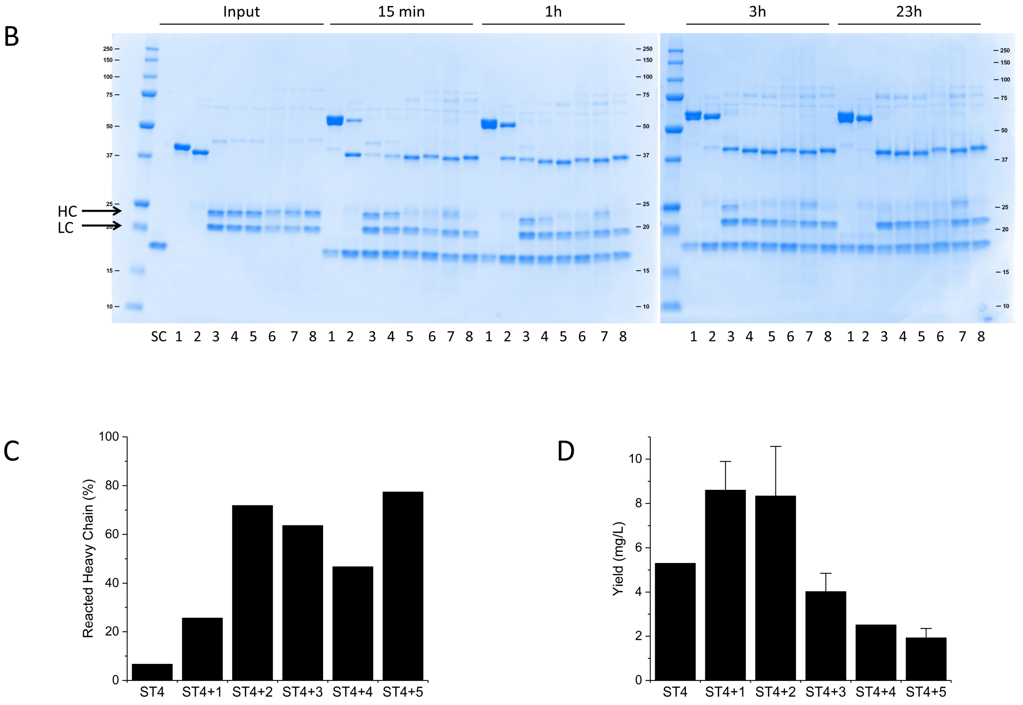


**Figure S5: Prevention of SpyTag Cleavage by Proximity to Folded Domains**

**A:** C-terminal sequences of 2 MBP constructs and 6 Fab heavy chains with a C-terminal SpyTag and varying linker lengths. **B:** SDS-PAGE of SpyCatcher (SC), the 8 constructs described in subfigure A (marked as 1-8) and their coupling reactions after 15 minutes, 1 hour, 3 hours, and 23 hours. **C:** Quantification of the reacted heavy chain band after 15 minutes coupling reaction with SpyCatcher. Quantification was performed with Image Lab 6.1 (Bio-Rad). **D:** Production yield of the Fab constructs in the formats described in this figure in TG1 F- cells (1 to 13 purifications per construct) in mg protein per liter of *E. coli* culture. Yields are absolute yields as determined by A280, and below 2 mg/L mostly consist of impurities. Error bars are standard deviations.

Figure S6

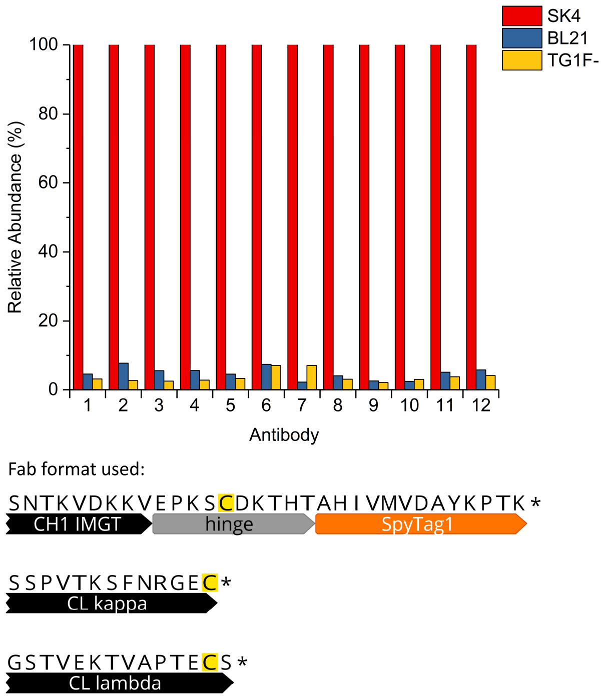


**Figure S6: Abundance of SpyTagged Fabs without Purification Tags in Bacterial Lysates**

Top: Relative abundance of full-length SpyTag containing Fabs for 12 different antibodies expressed in the format described by Alam et al.[1] measured in crude bacterial lysate from BL21 cells (used by Alam et al.), TG1 F- cells, and SK4 knockout cells. Relative abundance was determined by performing titration ELISAs with serial 1:2 dilutions of antibody containing bacterial lysate generated from an identical cell density (by OD600). ELISAs were performed by coating with 10 µg/mL SpyCatcher3 and detecting with anti-Fab-HRP. The ELISA curves were fitted by logistic regression and the EC50 values from the fits were calculated (Origin 2017, Origin labs). For each Fab, the lowest EC50 value in the sets of three cell types was chosen and divided by the individual EC50 values, resulting in 100% for the cell type with the highest abundance (and thus lowest EC50 value), plus the relative abundance for the other two cell types. In all cases, the concentration in lysate was at least a factor of 10 larger in SK4 cells than in BL21 or TG1 F-, suggesting that neither the presence of a heavy chain-light chain disulfide bond nor the absence of a purification tag has any influence on protease activity. Bottom: C-termini of the heavy and light chains of the Fab format used for these 12 binders. Cysteines are highlighted in yellow.

Figure S7

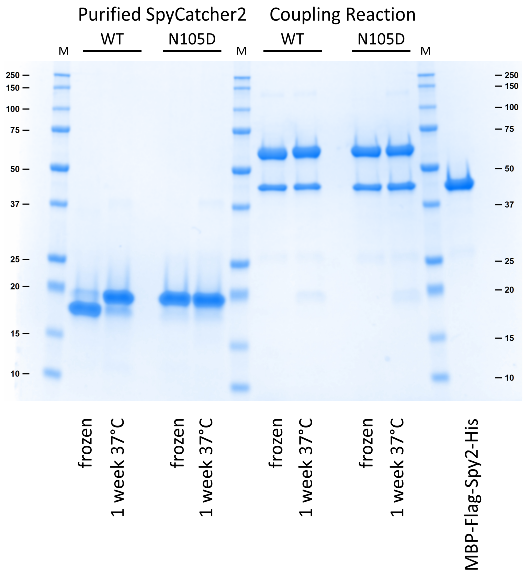


**Figure S7: Deamidation of SpyCatcher2**

SDS PAGE of SpyCatcher2 and SpyCatcher2 N105D mutant stored frozen and incubated for 1 week at 37°C and their coupling reactions with MBP-FLAG-SpyTag2-His. Incubation at 37°C causes a noticeable apparent molecular weight shift, which can be avoided by mutating asparagine 105 to aspartate. The coupling efficiency is not affected by the mutation or by incubation at 37°C. Marker lanes are indicated with ‘M’, molecular weights are given in kDa.

Figure S8

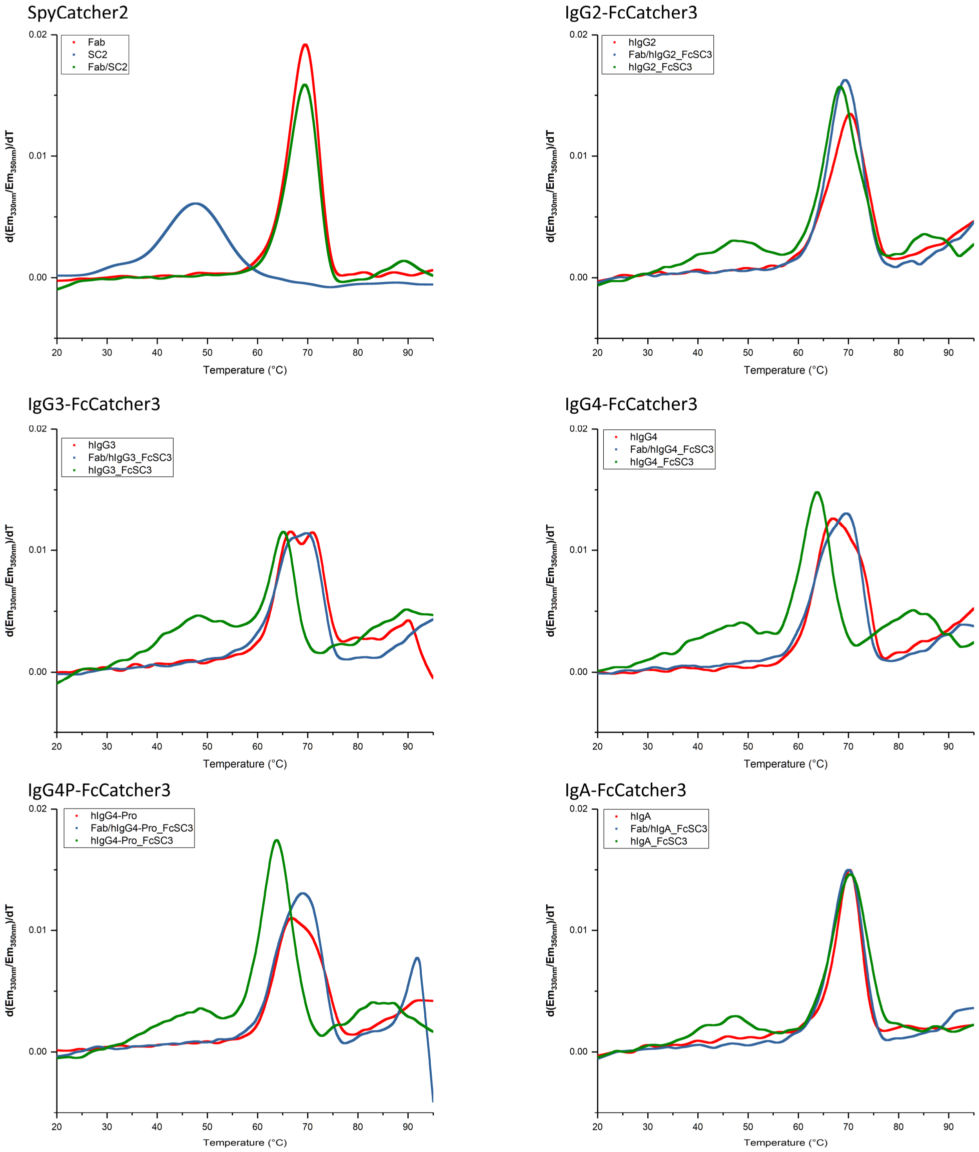


**Figure S8: Thermal Unfolding of SpyCatcher2 and FcCatcher Isotypes Before and After Coupling**

NanoDSF curves of SpyCatcher2, different human isotype FcCatchers and Fab+SpyCatcher/FcCatcher coupled products in comparison with the respective human isotype-matched full-length immunoglobulin or Fab. In all cases, a broad peak between 30 and 55°C is associated with uncoupled SpyCatcher and is not present in the coupled product. For some isotypes (IgG3, IgG4, IgG4-Pro), the unfolding of the SpyCatcher domain seems to lower the temperature of the main peak, corresponding to the unfolding of the IgG domains. This is not observed with coupled Fc-Catchers. Measurements were performed at approximately 0.4 mg/mL total protein concentration, except for SpyCatcher2, where measurements were performed at a concentration of 3 µM of Fab/SpyCatcher2.

Figure S9

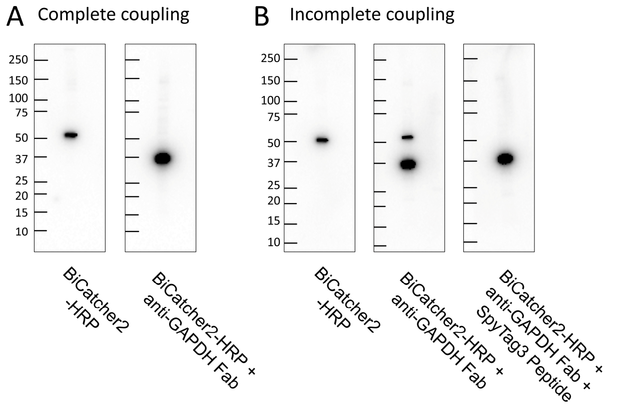


**Figure S9: Unspecific Binding of BiCatcher2-HRP on HeLa Cell Lysate in Immunoblots**

**A**: HeLa whole cell lysate (Bio-Rad) was blotted and probed via BiCatcher2-HRP or BiCatcher2-HRP coupled with an excess of anti-GAPDH Fab in FLAG-SpyTag2-His format. The 51 kDa band recognized by BiCatcher2-HRP alone is not recognized anymore after complete coupling with the Fab, instead a band with the molecular weight of GAPDH (37 kDa) is recognized. Coupling concentration: 4.4 µM Fab, 1.75 µM BiCatcher-HRP **B**: Sub-stochiometric coupling (here: 0.5 SpyTagged Fabs per 1 SpyCatcher domain) leads to recognition of both the antigen and the additional band. This can be blocked by incubating the coupled product with a 5-fold molar excess over BiCatcher2-HRP of SpyTag3 peptide. Coupling conditions: 1.75 µM Fab, 1.75 µM BiCatcher-HRP, 8.75 µM SpyTag3 peptide.

**Supplementary Table 1: Primers Used For Knockout Strain Generation**

| Oligo ID | Sequence |
|---|---|
| 119_SKE | CCTGGTGTTCTGAAACGGAGGCCGGGCCAGGCATGAACATGTTTTTTAGGGTGTAGGCTGGA GCTGCTTC |
| 120_SKE | TTTTCAAGCTTCGCCAGATCGAGTGCGATATTCACCGTCTCATCCAGATAATTCCGGGGATCCG TCGACC |
| 88_SKE | TCAGCTCTGACTGTCGGACA |
| 89_SKE | GACCATTACGGCCAGGTTC |
| 127_SKE | TAAAAAATACATATTCAATCATTAAAACGATTGAATGGAGAACTTTTATGTGTAGGCTGGAGCT GCTTCG |
| 128_SKE | CCTCGGGGAAATTTTAGTTGGCGTTCTTAAAATGTGTACTTAAGACCAGCATTCCGGGGATCCG TCGACC |
| 135_SKE | CGCTTATATGCCTCTAAAGG |
| 136_SKE | ATAATCGGCGTCACAGATG |

### Supplementary Text 1: Protein Sequences

**Fab Formats**

In all light chains, the final cysteine is removed. For light chain, the ompA signal peptide (MKKTAIAIAVALAGFATVAQA) was used. For heavy chains, the phoA signal peptide (MKQSTIALALLPLLFTPVTKA) was used.

Legend: SpyTag Linker Hinge FLAG-tag His-tag Signal Peptide

**Human Fab heavy chains:**

Flag-SpyTag1-His

VH-CH1…TKVDKKV-EPKS-EF-DYKDDDDK-GGS-AHIVMVDAYKPTK-GAP-HHHHHH

His-SpyTag1-Flag

VH-CH1… TKVDKKV-EPKS-EF-HHHHHH-GAPG-AHIVMVDAYKPTK-GGS-DYKDDDDK

Flag-SpyTag2-His

VH-CH1… TKVDKKV-EPKS-EF-DYKDDDDK-GGS-VPTIVMVDAYKRYK-GAP-HHHHHH

Flag-SpyTag3-His

VH-CH1… TKVDKKV-EPKS-EF-DYKDDDDK-GGS-RGVPHIVMVDAYKRYK-GAP-HHHHHH

ST4+2 SpyTag1-His

VH-CH1…TKVDKKV-EPKS-AHIVMVDAYKPTK-GAP-HHHHHH

**Murine Fab heavy chains**

Flag-SpyTag2-His

VH-CH1… TKVDKKI-EPRG-EF-DYKDDDDK-GGS-VPTIVMVDAYKRYK-GAP-HHHHHH

**Other spytagged proteins**

MBP-Flag-SpyTag-His

MKKTAIAIAVALAGFATVAQA-KIEEGKLVIWINGDKGYNGLAEVGKKFEKDTGIKVTVEHPDKLEEKFPQVAATGDGP
DIIFWAHDRFGGYAQSGLLAEITPDKAFQDKLYPFTWDAVRYNGKLIAYPIAVEALSLIYNKDLLPNPPKTWEEIPALDKE
LKAKGKSALMFNLQEPYFTWPLIAADGGYAFKYENGKYDIKDVGVDNAGAKAGLTFLVDLIKNKHMNADTDYSIAEAA

FNKGETAMTINGPWAWSNIDTSKVNYGVTVLPTFKGQPSKPFVGVLSAGINAASPNKELAKEFLENYLLTDEGLEAVNK
DKPLGAVALKSYEEELAKDPRIAATMENAQKGEIMPNIPQMSAFWYAVRTAVINAASGRQTVDEALKDAQT-EF-DYK
DDDDK-GGS-AHIVMVDAYKPTK-GAP-HHHHHH

MBP-His-SpyTag-Flag

MKKTAIAIAVALAGFATVAQA-KIEEGKLVIWINGDKGYNGLAEVGKKFEKDTGIKVTVEHPDKLEEKFPQVAATGDGP
DIIFWAHDRFGGYAQSGLLAEITPDKAFQDKLYPFTWDAVRYNGKLIAYPIAVEALSLIYNKDLLPNPPKTWEEIPALDKE
LKAKGKSALMFNLQEPYFTWPLIAADGGYAFKYENGKYDIKDVGVDNAGAKAGLTFLVDLIKNKHMNADTDYSIAEAA
FNKGETAMTINGPWAWSNIDTSKVNYGVTVLPTFKGQPSKPFVGVLSAGINAASPNKELAKEFLENYLLTDEGLEAVNK
DKPLGAVALKSYEEELAKDPRIAATMENAQKGEIMPNIPQMSAFWYAVRTAVINAASGRQTVDEALKDAQT-EF-HHH
HHH-GAPG-AHIVMVDAYKPTK-GGS-DYKDDDDK

ST4 MBP-SpyTag-His

MKKTAIAIAVALAGFATVAQA-KIEEGKLVIWINGDKGYNGLAEVGKKFEKDTGIKVTVEHPDKLEEKFPQVAATGDGP
DIIFWAHDRFGGYAQSGLLAEITPDKAFQDKLYPFTWDAVRYNGKLIAYPIAVEALSLIYNKDLLPNPPKTWEEIPALDKE
LKAKGKSALMFNLQEPYFTWPLIAADGGYAFKYENGKYDIKDVGVDNAGAKAGLTFLVDLIKNKHMNADTDYSIAEAA
FNKGETAMTINGPWAWSNIDTSKVNYGVTVLPTFKGQPSKPFVGVLSAGINAASPNKELAKEFLENYLLTDEGLEAVNK
DKPLGAVALKSYEEELAKDPRIAATMENAQKGEIMPNIPQMSAFWYAVRTAVINAASGRQTVDEALKDAQT- AHIVM
VDAYKPTK-GAP- HHHHHH

MBP-Flag-SpyTag-His (cytoplasmic)

MV-KIEEGKLVIWINGDKGYNGLAEVGKKFEKDTGIKVTVEHPDKLEEKFPQVAATGDGPDIIFWAHDRFGGYAQSGLL
AEITPDKAFQDKLYPFTWDAVRYNGKLIAYPIAVEALSLIYNKDLLPNPPKTWEEIPALDKELKAKGKSALMFNLQEPYFT
WPLIAADGGYAFKYENGKYDIKDVGVDNAGAKAGLTFLVDLIKNKHMNADTDYSIAEAAFNKGETAMTINGPWAWS
NIDTSKVNYGVTVLPTFKGQPSKPFVGVLSAGINAASPNKELAKEFLENYLLTDEGLEAVNKDKPLGAVALKSYEEELAKD
PRIAATMENAQKGEIMPNIPQMSAFWYAVRTAVINAASGRQTVDEALKDAQT-EF-DYKDDDDK-GGS-
AHIVMVDAYKPTK-GAP-HHHHHH

**Supplementary References**

1. Alam MK, Gonzalez C, Hill W, Fonge H, Barreto K, Geyer CR. Synthetic Modular Antibody Construction Using the SpyTag/SpyCatcher Protein Ligase System. *Chembiochem*, (2017).
